# Emergence of consciousness and complexity amidst diffuse delta rhythms: the paradox of Angelman syndrome

**DOI:** 10.1101/697862

**Authors:** Joel Frohlich, Lynne M. Bird, John Dell’Italia, Micah A. Johnson, Joerg F. Hipp, Martin M. Monti

## Abstract

Numerous theories link consciousness to informationally rich, complex neural dynamics. This idea is challenged by the observation that children with Angelman syndrome (AS), while fully conscious, display a hypersynchronous electroencephalogram (EEG) phenotype typical of information-poor dynamics associated with unconsciousness. If informational complexity theories are correct, then sufficiently complex dynamics must still exist during wakefulness and exceed that observed in sleep despite pathological delta (1 – 4 Hz) rhythms in children with AS. As characterized by multiscale metrics, EEGs from 35 children with AS feature significantly greater complexity during wakefulness compared with sleep, even when comparing the most pathological segments of wakeful EEG to the segments of sleep EEG least likely to contain conscious experiences, and when factoring out delta power differences across states. These findings support theories linking consciousness with complexity and warn against reverse inferring an absence of consciousness solely on the basis of clinical readings of EEG.

## Introduction

Electroencephalography (EEG) offers a window into neural activity during sleep and wakefulness, generally revealing low voltage, fast activity during wakefulness and high voltage, slow activity during non-rapid eye movement (NREM) sleep^1–4^. The former may be conceptualized as neural “chatter,” i.e., informationally rich interactions analogous to background noise at a sports arena, whereas the latter may be conceptualized as neural “chanting,” i.e., informationally poor synchronization analogous to coordinated crowd activity^5^. Similarly to NREM sleep, loss of consciousness in other states also coincides with a chanting EEG rhythm^6–8^ reflected in lower signal complexity^4, 9–13^. For instance, in a state of anesthesia, loss of consciousness coincides with a widespread increase in EEG power at low frequencies^3, 14, 15^, marking a decrease in corticocortical interactions^16, 17^. Absence seizures and temporal lobe seizures that impair consciousness are also associated with significant increases in slow waves^18, 19^. Similar findings have been reported for other modes of loss of consciousness including advanced states of encephalopathy and coma^20, 21^, sudden acceleration^22, 23^, basilar artery migraine^24^, and convulsive syncope^25^.

The association between neural chanting and loss of consciousness is consistent with the integrated information theory of consciousness (IIT)^26, 27^, which asserts that spatially extended, low-frequency (i.e., delta) rhythms lead to a loss of information differentiation^28^ and a breakdown of effective connectivity within the thalamocortical system^29^, as neural signaling pauses diffusely at the troughs of delta oscillations^30, 31^ and loss of consciousness results. In translational work, consciousness is also linked to neural complexity by the perturbational complexity index, a successful method of inferring consciousness based on the brain’s electrophysiological “echo” following a magnetic pulse^9, 32^.

In apparent contradiction to the above arguments and data, children with Angelman syndrome (AS) display the rich spectrum of purposeful behavior that implies conscious awareness ([as seen here](https://youtu.be/U5J0kvFSTtA))^33–35^ while exhibiting the chanting EEG phenotype typical of states of reduced consciousness (Fig. 1)^36–39^. Although the AS EEG phenotype has long been described in clinical reports^40^, we are the first to characterize the degree to which the awake EEG in children with AS can support complex dynamics and, moreover, that these dynamics are demonstrably lowered as consciousness decreases during sleep.

**Figure 1.**
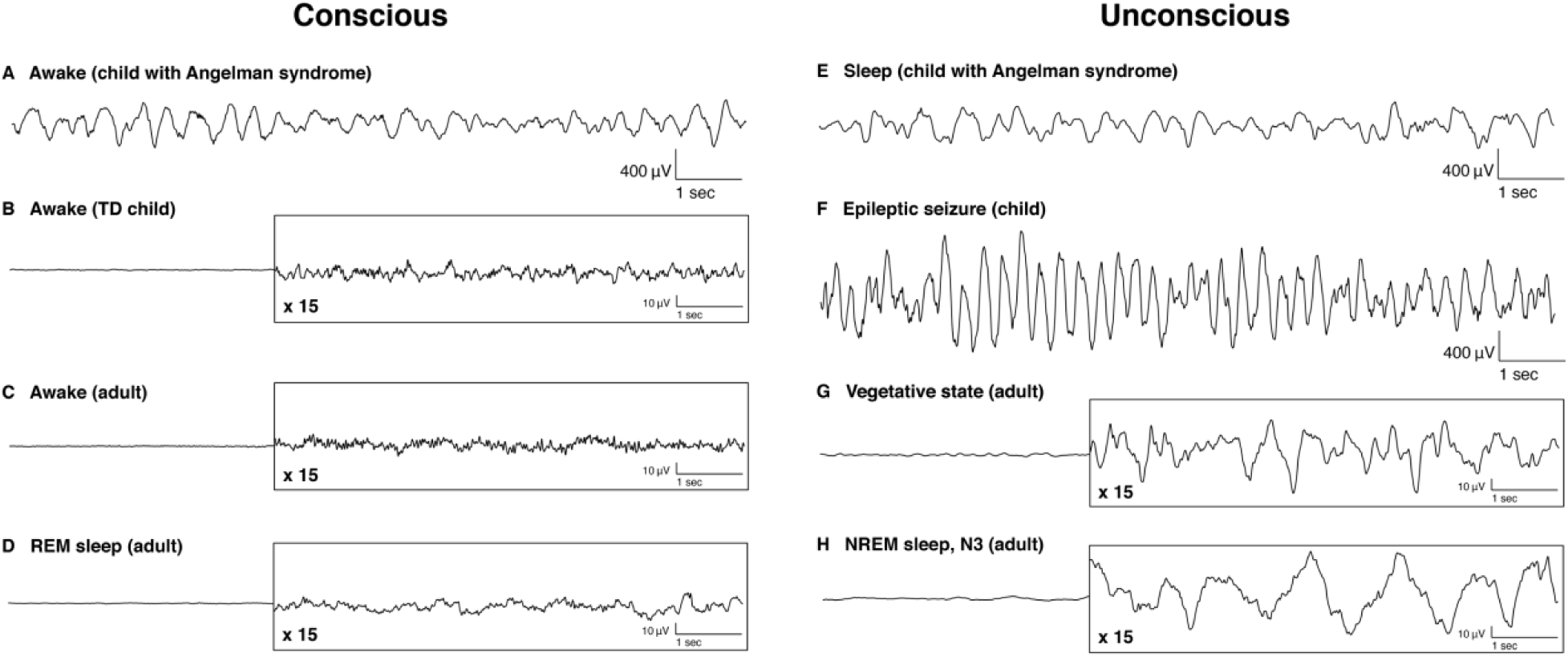
10 s EEG recordings from conscious states (left column) and unconscious states (right column). A general trend of low-voltage, fast activity is visible in all cases from conscious states except for AS, which paradoxically resembles unconscious EEG activity even during wakeful consciousness (panel A). Panels A and E display EEG from a 27-month-old girl with AS included in this study; other panels display data from outside this study with different acquisition, referencing, and preprocessing and are provided for illustrative purposes only. Direct, quantitative comparisons are precluded by these differences. Some EEGs are depicted with 15x amplitude exaggeration to better display waveforms (panels B,C,D,G,H). **(A)** Awake state EEG (channel Cz) from a 27-month-old girl with AS (Participant 10, Table S1) marked by high amplitude delta oscillations that are more typical of diminished consciousness (cf. left column). This participant did not have seizures and was not taking medication. **(B)** Awake state EEG (channel Cz) recorded from a typically developing 38-month old girl. **(C)** Awake state EEG (bipolar channel F1-F3) recorded from a healthy 37-year-old woman. **(D)** REM sleep EEG (bipolar channel F1-F3) recorded from a healthy 37-year-old woman. **(E)** Sleep EEG from a 27-old-girl with AS; note the extreme similarity in waveform to awake state AS EEG in A. **(F)** Ictal EEG (bipolar channel F3-C3) recorded from a 2-year-old girl with epilepsy during a seizure. **(G)** Spontaneous EEG (channel Cz) recorded from a 59-year-old man in a vegetative state. **(H)** Sleep EEG (bipolar channel F1-F3) recorded during NREM (stage N3) from a healthy 37-year-old woman.

**Table 1.** Channel-frequency (power) and channel-timescale (complexity) clusters identified using permutation cluster statistics. Bold rows are clusters reported in the text and figures that meet statistical significance after a Bonferroni correction (α = 0.0063). P-values are derived from empirical cluster size distributions using permutation tests. Effect sizes are reported as Cohen’s d (median and standard deviation across all cluster points; standard deviation is reported as N/A for clusters with only 1 point).

|  | EEG measure | Comparison | Direction (awake-sleep) | Regressed covariates? | p-value | Cohen's d (cluster median) | Cohen's d (cluster SD) | Cluster size | percentage | Low freq | High freq | Minimum number of channels | Maximum number of channels |
| --- | --- | --- | --- | --- | --- | --- | --- | --- | --- | --- | --- | --- | --- |
| **Cluster 1** | **gMLZ** | **Full** | **Increase** | **No** | **< 10<sup>-4</sup>** | **0.63** | **0.32** | **345** | **90.79** | **1.0** | **28.6** | **9** | **19** |
| **Cluster 2** | **power** | **Full** | **Decrease** | **No** | **< 10<sup>-4</sup>** | **-0.48** | **0.26** | **486** | **62.39** | **1.0** | **20.7** | **1** | **19** |
| **Cluster 3** | **mMSE** | **Full** | **Increase** | **No** | **0.0007** | **0.69** | **0.12** | **231** | **60.79** | **6.3** | **100.0** | **1** | **19** |
| **Cluster 4** | **power** | **Full** | **Increase** | **No** | **0.017** | **0.56** | **0.26** | **83** | **10.65** | **19.0** | **32.0** | **3** | **19** |
| **Cluster 5** | **gMLZ** | **Targeted** | **Increase** | **No** | **< 10<sup>-4</sup>** | **0.64** | **0.32** | **279** | **73.42** | **1.0** | **28.6** | **1** | **19** |
| **Cluster 6** | **mMSE** | **Targeted** | **Increase** | **No** | **0.001** | **0.67** | **0.11** | **226** | **59.47** | **6.3** | **100.0** | **3** | **19** |
| **Cluster 7** | **power** | **Targeted** | **Decrease** | **No** | **0.0011** | **-0.60** | **0.19** | **177** | **22.72** | **1.0** | **2.2** | **8** | **19** |
| **Cluster 8** | **power** | **Targeted** | **Increase** | **No** | **0.0022** | **0.63** | **0.24** | **157** | **20.15** | **11.3** | **32.0** | **2** | **19** |
| **Cluster 9** | **power** | **Targeted** | **Increase** | **No** | **0.1727** | **0.24** | **0.02** | **4** | **0.51** | **4.8** | **6.2** | **1** | **1** |
| **Cluster 10** | **power** | **Targeted** | **Increase** | **No** | **0.2021** | **0.27** | **0.02** | **3** | **0.39** | **4.8** | **5.7** | **1** | **1** |
| **Cluster 11** | **power** | **Targeted** | **Increase** | **No** | **0.2448** | **0.25** | **0.05** | **2** | **0.26** | **5.2** | **5.2** | **2** | **2** |
| **Cluster 12** | **power** | **Targeted** | **Increase** | **No** | **0.3036** | **0.24** | **N/A** | **1** | **0.13** | **4.0** | **4.0** | **1** | **1** |
| **Cluster 13** | **gMLZ** | **Targeted** | **Increase** | **Yes** | **< 10<sup>-4</sup>** | **2.42** | **1.28** | **361** | **95.00** | **1.0** | **28.6** | **10** | **19** |
| **Cluster 14** | **mMSE** | **Targeted** | **Increase** | **Yes** | **< 10<sup>-4</sup>** | **1.92** | **0.81** | **346** | **91.05** | **5.0** | **100.0** | **11** | **19** |
| **Cluster 15** | **gMLZ** | **Targeted** | **Decrease** | **Yes** | **0.0844** | **-0.92** | **N/A** | **1** | **0.26** | **1.0** | **1.0** | **1** | **1** |
| **Cluster 16** | **gMLZ** | **Targeted** | **Decrease** | **Yes** | **0.0844** | **-0.58** | **N/A** | **1** | **0.26** | **1.0** | **1.0** | **1** | **1** |
| **Cluster 17** | **gMLZ** | **Targeted** | **Decrease** | **Yes** | **0.0844** | **-0.71** | **N/A** | **1** | **0.26** | **1.0** | **1.0** | **1** | **1** |

AS is caused by dysfunction of the gene *UBE3A*^41, 42^. Its clinical phenotype encompasses global developmental delay, intellectual disability, microcephaly, epilepsy, and sleep difficulties^42–44^. Puzzlingly, awake state EEG recordings from children with AS display diffuse, slow rhythmic oscillations at delta (1-4 Hz) frequencies^36–38^ reminiscent of those seen in slow wave sleep. In fact, spectral power at the delta peak frequency (2.8 Hz) in awake children with AS exceeds that observed in typically developing (TD) children by > 1000%^36^. At face value, either the core ideas of IIT do not generalize to the hypersynchronized, but wakeful and conscious, brain in AS, or amidst the pathologically slow chanting rhythm observed in awake children with AS, sufficiently complex interactions nonetheless arise and persist over time despite the periodic presence of the widespread interburst silence at the trough of each delta wave, generally assumed to interrupt the flow of consciousness^4, 28–30^.

To address this puzzle, we examined whether brain dynamics observed in children with AS during periods of wakefulness were measurably greater than those observed during periods of sleep, as predicted by complexity-based theories of consciousness and despite the diffuse presence of large delta oscillations in both states. As described below, contrary to common readings of EEG and despite diffuse delta oscillations, the awake EEG of children with AS supports significantly greater signal complexity than the asleep AS EEG. This finding persisted even after contrasting periods of wakefulness showing the most pathological EEG signature to the periods of sleep least likely to coincide with any oneiric experience^45, 46^, and, moreover, after accounting for differences in delta power between states.

## Results

Herein, we assess the degree to which signal complexity can emerge from the pathologically hypersynchronous brain dynamics typical of children with AS and, specifically, whether such dynamics differ significantly across levels of consciousness (i.e., wake, sleep). We first address this question by analyzing all useable EEG data (henceforth, full comparison). We then repeat the analysis while controlling for two possible confounds (henceforth, targeted comparison). The sample included a cohort of 35 children with AS (15 female), ranging from 13 to 130 months of age (mean ± std = 47.9 ± 28.6 months), of which 25 had a deletion of chromosome 15q11-q13 (see Table S1 for individual demographic details and length of EEG data used in each of the analyses). EEG recordings were acquired in a clinical setting using an international 10-20 EEG montage (19 channels), and sleep EEG data were collected and marked by a technician as children slept naturally during the EEG session. Consistent with prior studies of AS^36–39, 47^, qualitative inspection of EEG recordings revealed strongly abnormal EEG patterns in both sleep and wakefulness (see Fig. 1A,E for examples from a participant without seizures or medications).

To assess biomarkers for consciousness in AS, we derived EEG spectral power using a Morlet wavelet transform and EEG complexity using two measures, modified multiscale entropy (mMSE) and generalized multiscale Lempel-Ziv (gMLZ). mMSE captures the balance between periodicity and randomness in the signal, computed as modified sample entropy (mSampEn)^48^ across 20 time-scales using a coarse graining procedure that excludes high frequencies at each step^49^. gMLZ captures the incompressibility or number of unique substrings in the signal^50^ and is also computed across 20 timescales^51^ using two median filters with different smoothing windows to exclude both low and high frequencies at each step^52^. We accounted for multiple comparisons using cluster randomization statistics^53, 54^ to identify clusters in channel-frequency (power) and channel-timescale (complexity) space that show significant changes with sleep (one p-value derived per cluster using permutation tests). We then adjusted for 8 separate tests (full comparison: 3 EEG measures, targeted comparison: 3 EEG measures, targeted comparison covarying for delta power, 2 EEG measures) using a Bonferroni correction, yielding a test-wise criterion of α = 0.0063.

## Full Comparison

The full comparison revealed peaks in the delta band for both the awake and asleep condition (channel-averaged), with a sharper peak in the awake state and a broader peak in the asleep state (Fig. 2A; see Fig. S1A for visualization of the untransformed power). The duration of useable EEG ranged from 3.39 to 167 minutes (awake state, mean ± std = 16.7 ± 27.1 minutes) and 2.89 to 123 minutes (asleep state, mean ± std = 17.3 ± 20.1 minutes). Power was generally decreased in wakefulness at frequencies under 20 Hz, with the largest decrease occurring as a 53.0% reduction at f = 1.52 Hz (Fig. 2B, see Fig. S1B for percent change referenced to wakefulness). These changes in wakeful power mapped onto a significant cluster (p < 10^−4^, cluster permutation test) in channel-frequency space with a spatially defuse topography that was largest over frontocentral areas (Fig. 2C,D; effect size: d = −0.48 ± 0.26, median ± SD; see Table 1 for full details of clusters). Because the spectral profile of this cluster appeared to “fuse” an oscillatory change in the delta frequency range with an oscillatory change in the alpha/beta frequency range (Fig. 2D), we then repeated the cluster randomization statistics with a stricter threshold (p = 0.0005) to observe the topography of each oscillatory change separately (Fig. S2, Table S2). We also observed a > 100% increase in power at high frequencies (f > 28 Hz) in the awake state relative to the asleep state (Fig. 2B); however, the corresponding cluster did not reach statistical significance (Cluster 4, Table 1).

**Figure 2.**
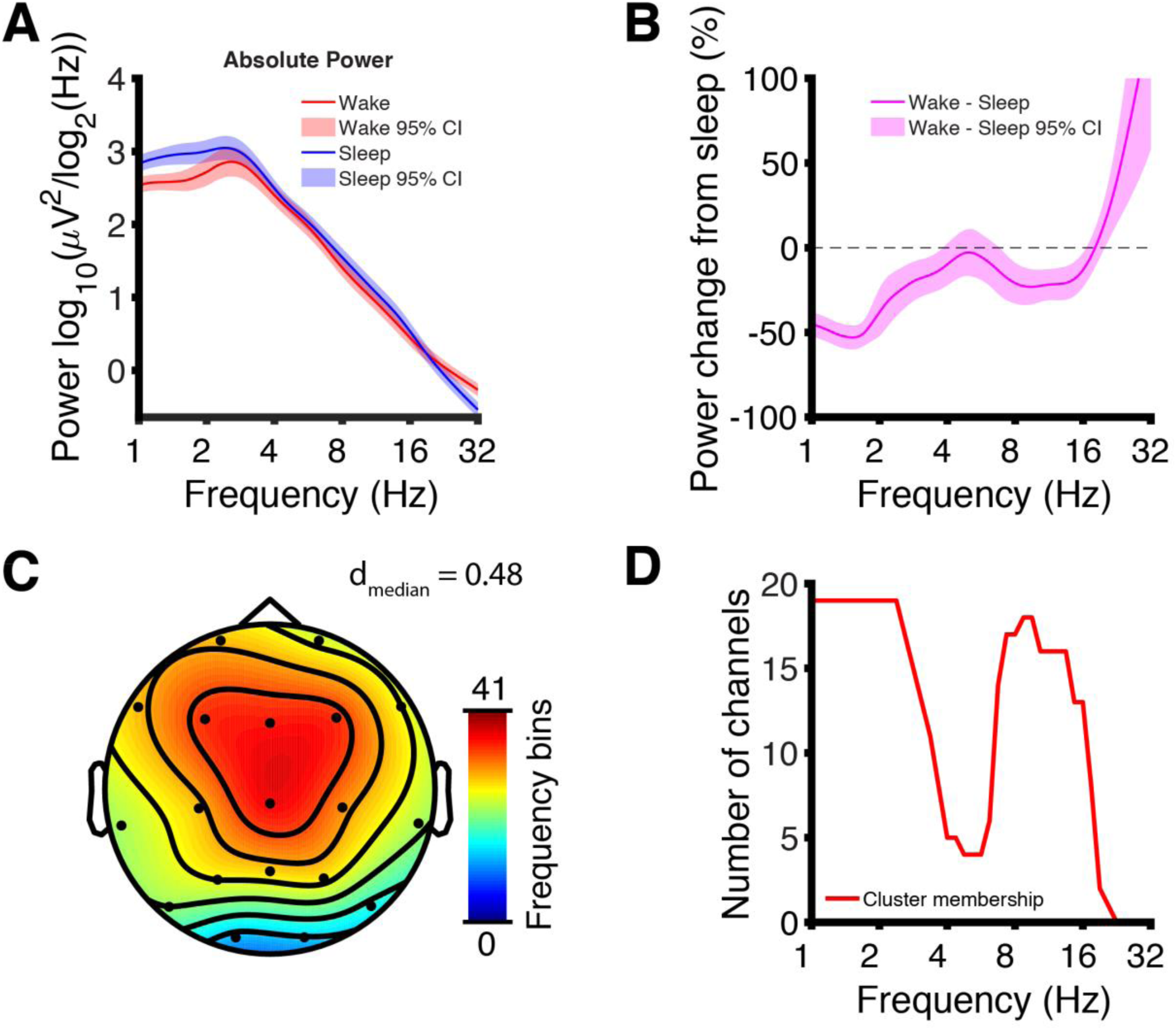
Full comparison of EEG power in sleep vs wakefulness. **(A)** Channel-averaged EEG power traces demonstrate global maxima in the delta band. **(B)** Percent power change from sleep after channel averaging; the largest change is a 53.0% decrease in power in wakefulness at 1.52 Hz. Increased power >20 Hz in wakefulness may indicate residual muscle artifact. **(C)** Channel-frequency cluster (p < 10^−4^, d_median_ = 0.48) of decreased power in wakefulness color-coded by the number of frequency bins participating in the cluster at each channel. The cluster was largest along its frequency dimension at channels Cz, Fz, F3, and F4. **(D)** Channel-frequency cluster membership plotted as the number of channels participating in the cluster at each frequency bin. The cluster was largest along its spatial dimension at delta and beta frequencies and smallest at theta and alpha frequencies.

**Table 2:** gMLZ timescale parameters. To compute Lempel-Ziv complexity (i.e., the difficulty of compressing in the signal), we first apply separately two median filters at each timescale: one filter with a smaller kernel (smoothing window) and a second filter with a larger kernel (thresholding window). The output from the first filter is then binarized according to the output from the second filter, which acts as a dynamic threshold. Lempel-Ziv complexity is then computed from the binary timeseries. Smoothing and thresholding windows are both spaced logarithmically to allow for good coverage of the EEG spectrum at all frequency bands, and the difference in size between the smoothing window and the thresholding window is varied to allow larger bandwidth at higher frequencies.

| *Scale* | **Center**<br>**frequency**<br>**(Hz)** | **Bandwidth**<br>**(Hz)** | **Threshold**<br>**frequency**<br>**(Hz)** | **Smoothing**<br>**frequency**<br>**(Hz)** | **Thresholding**<br>**window**<br>**(samples)** | **Smoothing**<br>**window**<br>**(samples)** |
| --- | --- | --- | --- | --- | --- | --- |
| 1 | 1.00 | 0.40 | 0.80 | 1.20 | 251 | 167 |
| 2 | 1.18 | 0.47 | 0.95 | 1.42 | 211 | 141 |
| 3 | 1.42 | 0.58 | 1.13 | 1.71 | 177 | 117 |
| 4 | 1.68 | 0.68 | 1.34 | 2.02 | 149 | 99 |
| 5 | 1.98 | 0.78 | 1.57 | 2.35 | 127 | 85 |
| 6 | 2.35 | 0.95 | 1.87 | 2.82 | 107 | 71 |
| 7 | 2.82 | 1.14 | 2.25 | 3.39 | 89 | 59 |
| 8 | 3.39 | 1.34 | 2.74 | 4.08 | 73 | 49 |
| 9 | 3.92 | 1.48 | 3.17 | 4.65 | 63 | 43 |
| 10 | 4.65 | 1.94 | 3.77 | 5.71 | 53 | 35 |
| 11 | 5.71 | 2.25 | 4.65 | 6.90 | 43 | 29 |
| 12 | 6.90 | 2.59 | 5.41 | 8.00 | 37 | 25 |
| 13 | 8.00 | 3.07 | 6.45 | 9.52 | 31 | 21 |
| 14 | 9.52 | 4.36 | 7.41 | 11.76 | 27 | 17 |
| 15 | 11.76 | 3.81 | 9.52 | 13.33 | 21 | 15 |
| 16 | 13.33 | 4.86 | 10.53 | 15.38 | 19 | 13 |
| 17 | 15.38 | 6.42 | 11.76 | 18.18 | 17 | 11 |
| 18 | 18.18 | 6.84 | 15.38 | 22.22 | 13 | 9 |
| 19 | 22.22 | 10.39 | 18.18 | 28.57 | 11 | 7 |
| 20 | 28.57 | 17.78 | 22.22 | 40.00 | 9 | 5 |

We next examined mMSE and gMLZ and found greater EEG signal complexity in the awake state as compared to sleep. Specifically, the channel averaged mSampEn decreased monotonically with faster timescales, but was greater during wakefulness as compared to sleep (Fig. 3A,B), with the exception of frequencies ≤ 6.25 Hz (i.e., the 5 slowest timescales). Greater mMSE during wakefulness was marked by a significant cluster (p = 0.0007) covering all channels but largest over central and posterior areas (Fig. 3C,D; effect size: d = 0.69 ± 0.12, median ± SD). By comparison, gMLZ increased monotonically with faster timescales and was larger in wakefulness as compared to sleep at all timescales, particularly those with center frequencies corresponding to delta and beta frequencies (channel-averaged; Fig. 3E,F). These changes were accompanied by a significant cluster (p < 10^−4^) encompassing 90.8% of channel-timescale space (Fig. 3G,H; effect size: d = 0.63 ± 0.32, median ± SD). The cluster appeared to fuse complexity changes corresponding to fast and slow timescales. We thus repeated the analysis with a stricter threshold (p = 0.0005) to view the topography of each change separately. Effect sizes in the low-frequency cluster were large (d = 0.96 ± 0.27; Fig. S3, Table S4). See Fig. S4 for complete visualizations of all clusters.

**Figure 3.**
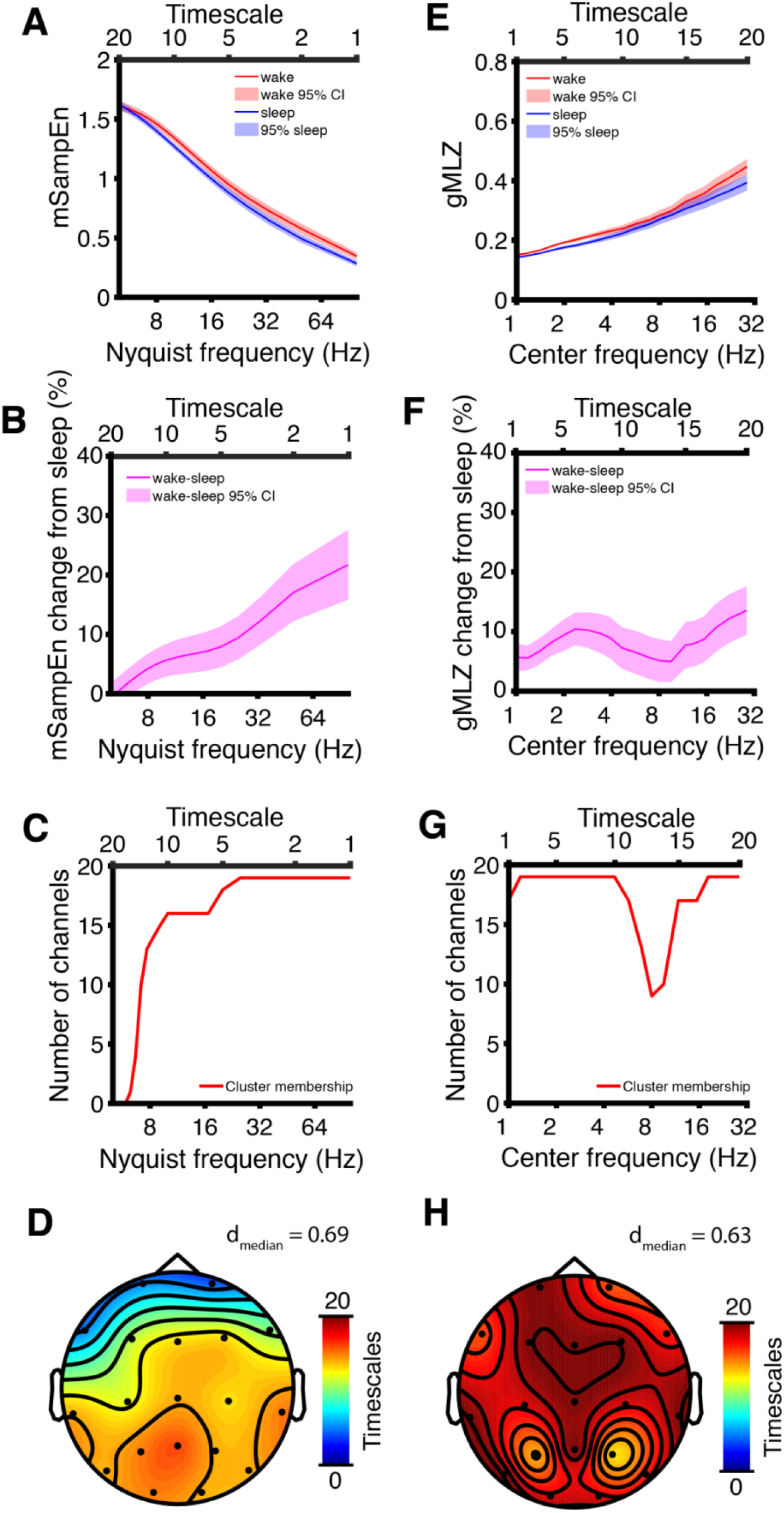
Full comparison of EEG complexity in sleep vs wakefulness. **(A)** Channel-averaged mMSE curves show greater complexity (i.e., mSampEn) at slower timescales (i.e., timescales with greater coarse graining). Nyquist frequency is indicated on the lower ordinate to indicate coarse graining, with the corresponding timescale given on the upper ordinate. **(B)** Percent change in channel-averaged mSampEn (wakefulness – sleep); the largest changes are observed at fast timescales (i.e., timescales with less coarse graining). **(C)** Membership for a channel-timescale cluster (p = 0.0007, d_median_ = 0.69) of increased mMSE in wakefulness plotted as the number of channels participating in the cluster at each timescale. The cluster was largest along its spatial dimension at faster timescales. **(D)** Channel-timescale cluster of increased mMSE in wakefulness color-coded by the number of timescales participating in the cluster at each channel. The cluster grew largest along its frequency dimension traveling posteriorly. **(E)** Channel-averaged gMLZ curves show greater complexity at faster timescales (i.e., timescales with shorter filter windows). Center frequency is indicated on the lower ordinate with the corresponding timescale given on the upper ordinate. **(F)** Percent change in channel-averaged gMLZ (wakefulness – sleep); complexity was greater in wakefulness, with the largest increases at timescales corresponding to delta and beta frequencies. **(G)** Membership for a channel-timescale cluster (p < 10^−4^, d_median_ = 0.63) of increased gMLZ in wakefulness plotted as the number of channels participating in the cluster at each timescale. The cluster was largest along its spatial dimension at timescales corresponding to delta and beta timescales. **(H)** Channel-timescale cluster of increased gMLZ in wakefulness color-coded by the number of timescales participating in the cluster at each channel. The cluster was nearly saturated in space, with local minima at P3 and P4.

## Targeted comparison

The full comparison (above) may have been influenced by two confounding factors. First, there is considerable variance is the delta amplitude during the awake state in AS^55^. It is thus conceivable that our findings from the full comparison would disappear when only the most pathological sections of awake EEG are considered. Secondly, it has been previously shown that some form of conscious experience can take place in up to 70% of NREM sleep in healthy volunteers^56^. Thus, it is also conceivable that our sleep data are “contaminated” by conscious mentation. We therefore performed the targeted comparison below to control for these confounds as follows. We addressed the first concern by focusing only on the most abnormal segments of conscious, wakeful EEG, as operationalized by delta power. We then addressed the second concern by contrasting the pathological wakeful EEG with the periods of sleep EEG least likely to correspond to any oneiric experience, as operationalized by the ratio of parietal delta power to high-frequency power^45, 46^, where higher values of this ratio correspond to a greater probability of unconsciousness. In other words, we compared the most abnormal segments of EEG still corresponding, nonetheless, to a state of wakeful consciousness, to the segments of sleep EEG least likely to correspond to any conscious experience. By using only these segments, we would expect the findings from the full comparison to disappear if EEG complexity is not always greater during wakefulness as compared to dreamless sleep, e.g., during bouts of especially high amplitude delta in wakefulness^55^. The duration of selected EEG ranged from 2.07 to 9.44 minutes (awake state, mean ± std = 3.96 ± 1.67 minutes) and 1.48 to 5.32 minutes (asleep state, mean ± std = 3.24 ± 1.02 minutes).

Consistent with the data selection criteria which optimize delta power, and closely replicating the results described in the full comparison, the targeted awake EEG sections were characterized by a prominent peak in the delta band, while the targeted asleep EEG sections were characterized by two delta band peaks at different octaves (channel-averaged; Fig. 4A), which are best visualized in the untransformed power (Fig. S5A); this suggests the presence of two separate oscillatory processes, one related to sleep and one related more specifically to AS pathology. Decreases in power between the two states were restricted to the delta band (max change: 41.3% decrease at f = 1.34 Hz), with > 100% increases also occurring at high frequencies (f > 25 Hz) (Fig. 4B, see Fig. S5B for percent change referenced to wakefulness). Permutation cluster statistics identified two small but significant clusters differing between the two states. The first cluster corresponded to decreased power at low delta (1.0 – 2.2 Hz) frequencies in the awake state (p = 0.0011) and lacked a distinct scalp topography (Fig. 4C,D; effect size: d = −0.60 ± 0.19, median ± SD). The second cluster corresponded mostly to increased power at mostly beta (11.3 – 32 Hz) frequencies in the awake state (p = 0.0022) and displayed a scalp topography suggestive of neck muscle artifact (Fig. 4E,F; effect size: d = 0.63 ± 0.24, median ± SD). These results show that, after accounting for the confounds that motivated our targeted comparison, the most reliable spectral differences between sleep and wakefulness were found at low delta frequencies. Note that the effect sizes of the differences in power across states were larger in the targeted comparison than in the full comparison despite matching data on more stringent criteria, likely as a result of the larger cluster in the full comparison fracturing into smaller clusters focused on the channel-frequency subspaces with the largest effects.

**Figure 4.**
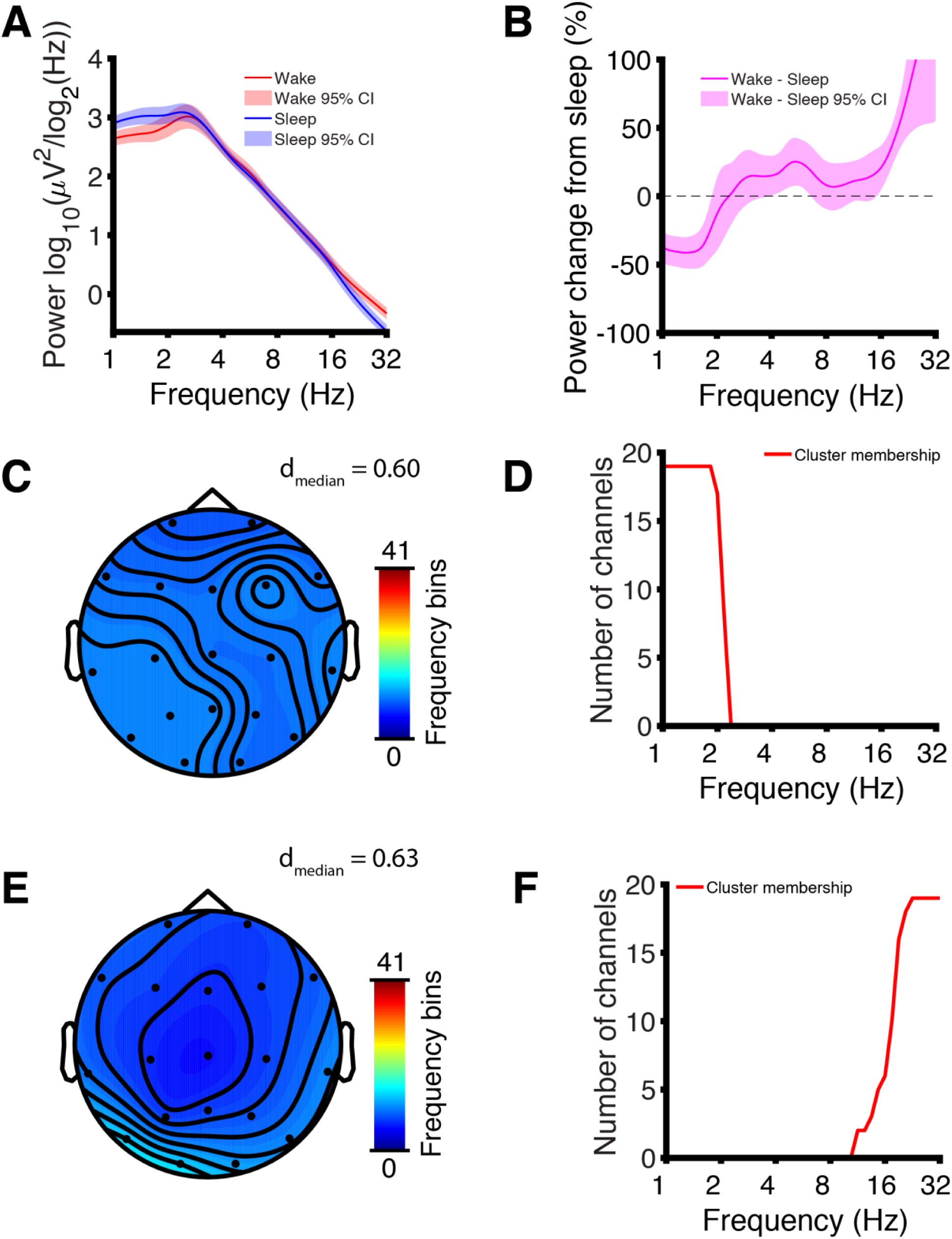
Targeted comparison of EEG power in sleep vs wakefulness. **(A)** Channel-averaged EEG power traces demonstrate global maxima in the delta band. **(B)** Percent power change from sleep after channel averaging; the largest change is a 41.3% decrease in power during wakefulness at 1.34 Hz. Increased power > 10.3 Hz in wakefulness may indicate either cortical arousal or muscle artifact. **(C)** Channel-frequency cluster (p = 0.0011, d_median_ = 0.60) of decreased power in wakefulness color-coded by the number of frequency bins participating in the cluster at each channel. All channels contributed roughly equally. **(D)** Channel-frequency cluster membership plotted as the number of channels participating in the cluster at each frequency bin. The cluster was saturated along its spatial extent at frequencies ≤ 1.8 Hz. **(E)** Channel-frequency cluster (p = 0.0022, d_median_ = 0.63) of increased power in wakefulness color-coded by the number of frequency bins participating in the cluster at each channel. The cluster was largest along its frequency dimension at channels T5 and O1 and displayed an overall topography suggestive of neck muscle artifact. **(F)** Channel-frequency cluster membership plotted as the number of channels participating in the cluster at each frequency bin. The cluster became saturated along its spatial extent at frequencies ≥ 22.6 Hz, suggesting possible involvement of residual muscle artifact in wakefulness.

Analysis of EEG signal complexity in the targeted comparison revealed similar results to the full comparison (channel-averaged; Fig. 5A,B). Greater mMSE in wakefulness was marked by a significant cluster (p = 0.001) exhibiting similar topography to the corresponding cluster found in the full comparison (Fig. 5C,D; effect size: d = 0.67 ± 0.11, median ± SD). The gMLZ curves also exhibited the same behavior seen in the full comparison (channel-averaged; Fig. 5E,F) and yielded a significant cluster (p < 10^−4^) marking greater complexity during wakefulness (Fig. 5G,H; effect size: d = 0.64 ± 0.32, median ± SD). The cluster appeared to fuse complexity changes corresponding to fast and slow timescales. We thus repeated the analysis with a stricter threshold (p = 0.0005) to view the topography of each change separately. Effect sizes in the low-frequency cluster were large (d = 0.98 ± 0.20, median ± SD; Fig. S6, Table S2). See Fig. S7 for complete visualizations of all clusters.

**Figure 5.**
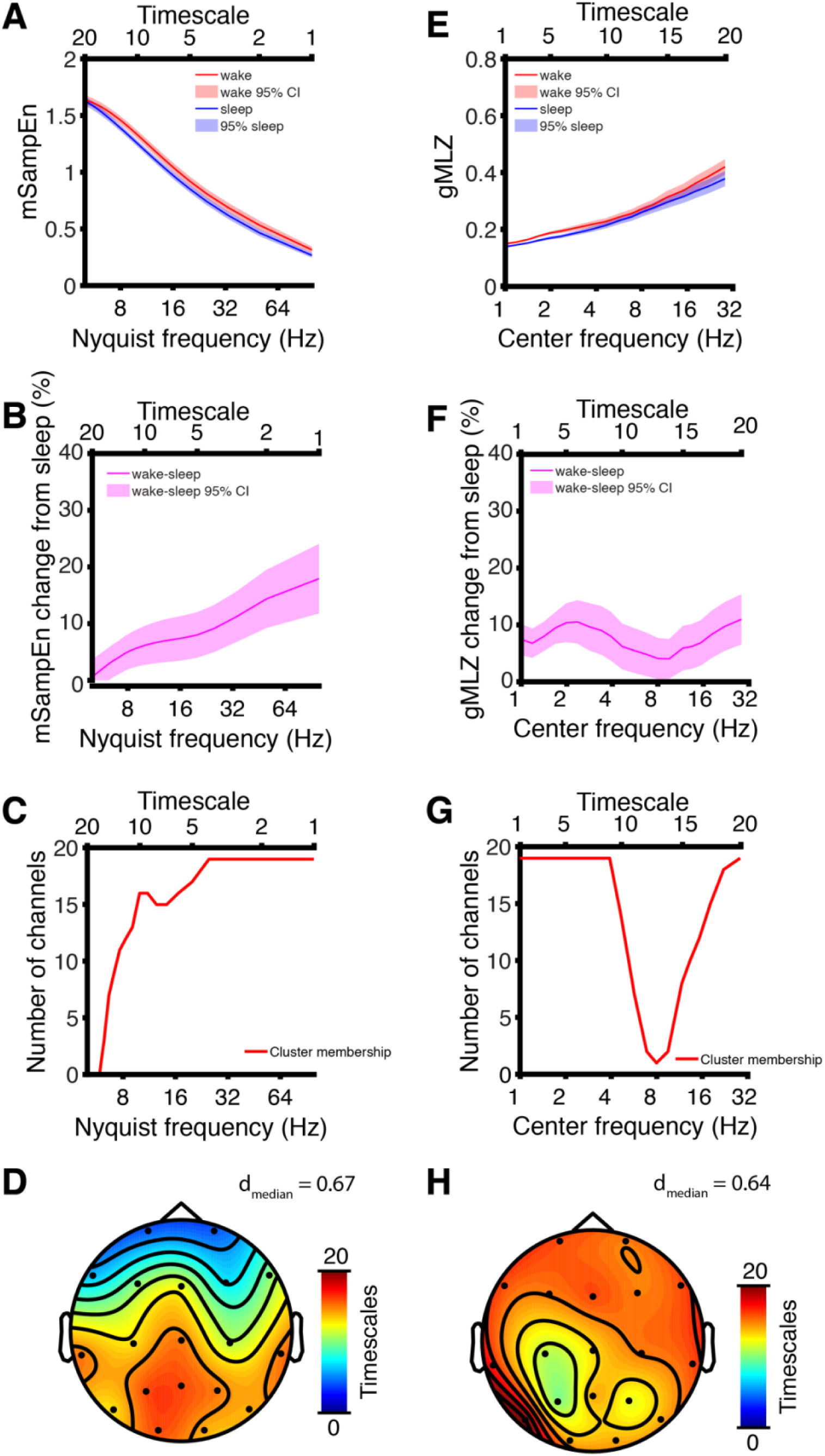
Targeted comparison of EEG complexity in sleep vs wakefulness. **(A)** Channel-averaged mMSE curves show greater complexity (i.e., mSampEn) at slower timescales (i.e., timescales with greater coarse graining). Nyquist frequency is indicated on the lower ordinate to indicate coarse graining, with the corresponding timescale given on the upper ordinate. **(B)** Percent change in channel-averaged mSampEn (wakefulness – sleep); the largest changes are observed at fast timescales (i.e., timescales with less coarse graining). **(C)** Membership for a channel-timescale cluster of increased mMSE in wakefulness plotted as the number of channels participating in the cluster at each timescale. The cluster was largest along its spatial dimension at faster timescales (channel saturation occurred at Nyquist frequencies ≥ 24.3 Hz). **(D)** Channel-timescale cluster (p = 0.001, d_median_ = 0.67) of increased mMSE in wakefulness color-coded by the number of timescales participating in the cluster at each channel. The cluster grew largest along its frequency dimension traveling posteriorly. **(E)** Channel-averaged gMLZ curves show greater complexity at faster timescales (i.e., timescales with shorter filter windows). Center frequency is indicated on the lower ordinate with the corresponding timescale given on the upper ordinate. **(F)** Percent change in channel-averaged gMLZ (wakefulness – sleep); complexity was greater in wakefulness, with the largest increases at timescales corresponding to delta and beta frequencies. **(G)** Membership for a channel-timescale cluster (p < 10^−4^, d_median_ = 0.64) of increased gMLZ in wakefulness plotted as the number of channels participating in the cluster at each timescale. The cluster was largest along its spatial dimension at timescales corresponding to delta and beta timescales (channel saturation occurred at timescales with center frequencies ≤ 3.9 Hz and = 28.6 Hz). **(H)** Channel-timescale cluster of increased gMLZ in wakefulness color-coded by the number of timescales participating in the cluster at each channel. The cluster was smallest in its frequency extent over parietocentral regions and nearly saturated elsewhere.

Finally, as shown in Fig. 4A, despite having selected the most pathological segments of the awake EEG dataset, delta power still differed significantly across wakefulness and sleep. Given the sinusoidal nature of delta oscillations, it is initially conceivable that the observed difference in complexity across the two states is nothing more than a trivial reflection of differences in delta power, i.e., high amplitude oscillations introducing strong regularities in the signal recorded at the scalp that diminish its complexity. Consistent with this view, delta power was negatively correlated with mSampEn at most timescales after averaging across channels, explaining the majority of the variance in mSampEn at fast timescales (R^2^ > 0.5 for Nyquist frequency ≥ 20 Hz, awake, and Nyquist frequency ≥ 50 Hz, asleep, Fig. 6A). Nonetheless, delta power did not mediate the effect of state (i.e., wakefulness vs sleep) on mMSE (Fig. 6B, p > 0.05 all timescales, uncorrected). Even more so than mMSE, delta power was negatively correlated with gMLZ at most timescales (R^2^ > 0.5 for center frequency ≥ 2.82 Hz, asleep and awake state, Fig. 6C). Yet again, delta power did not mediate the effect of state on gMLZ (Fig. 6D, p > 0.05 all timescales, uncorrected). Given the observed negative relationship between delta power and complexity measures, in what follows we repeated the targeted comparison after covarying for delta power (integrated 1 – 4 Hz).

**Figure 6.**
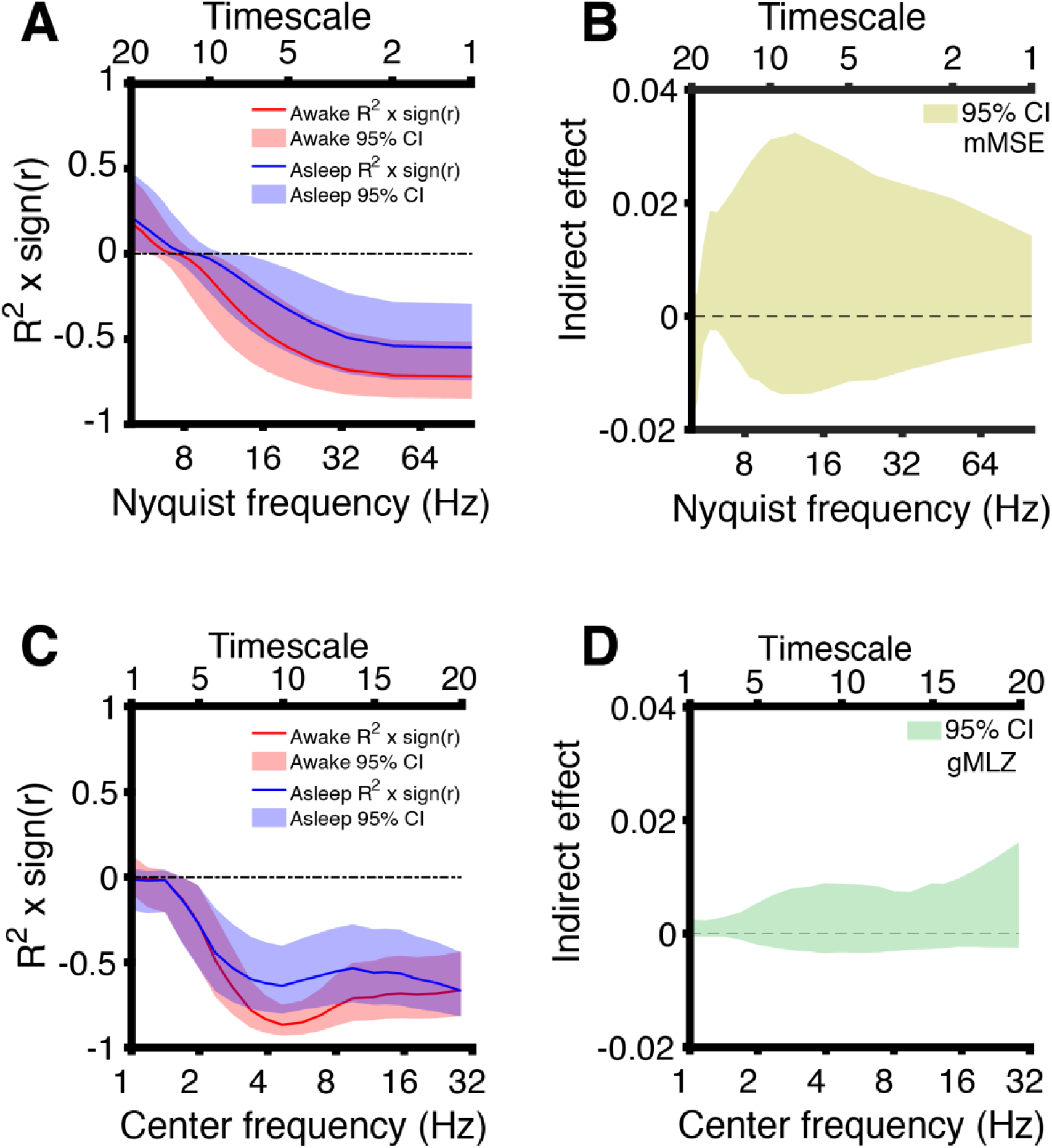
Targeted comparison of EEG complexity in sleep vs wakefulness after covarying for delta power. **(A)** Correlations (Pearson’s coefficient) between mSampEn and delta power computed after averaging across channels. Nyquist frequency is indicated on the lower ordinate to indicate coarse graining, with the corresponding timescale given on the upper ordinate. **(B)** Bootstrapped 95% confidence intervals of the indirect effect of sleep on mMSE mediated by delta power. Delta power did not mediate effects of sleep on mMSE at any timescale, even before correcting for multiple comparisons (p > 0.05 all timescales). **(C)** Correlations (Pearson’s coefficient) between gMLZ and delta power computed after averaging across channels. Center frequency is indicated on the lower ordinate with the corresponding timescale given on the upper ordinate**. (D)** Bootstrapped 95% confidence intervals of the indirect effect of sleep on gMLZ mediated by delta power. Delta power did not mediate effects of sleep on gMLZ at any timescale, even before correcting for multiple comparisons (p > 0.05 all timescales).

As shown in Fig. 7, our overall findings remained unchanged after controlling for delta power differences across wakefulness and sleep. Specifically, while mMSE curves were no longer monotonic with timescale, they still show the awake EEG to be more complex than the asleep EEG. The largest percent increase from sleep occurred at low frequencies (i.e., the fastest timescale; channel-averaged; see Fig. 7A,B). This relative increase in mMSE during wakefulness corresponded to a significant (p < 10^−4^) and nearly saturated cluster (Fig. 7 C,D; effect size: d = 1.92 ± 0.81, median ± SD). With respect to gMLZ, covarying for delta power again leads to the same qualitative result reported above, with wakefulness showing consistently greater complexity than sleep. Intriguingly, however, the divergence in complexity between the two states is even greater after factoring out delta power, with the largest percent increase from sleep (30.7%) occurring at the center frequency of 3.4 Hz (i.e., the 8^th^ timescale; channel-averaged; see Fig. 7 E,F). The relative increase in gMLZ during wakefulness corresponds to a significant (p < 10^−4^) and, again, nearly saturated cluster (Fig. 7 G,H; effect size: d = 2.42 ± 1.28, median ± SD). See Fig. S8 for complete visualizations of all clusters.

**Figure 7.**
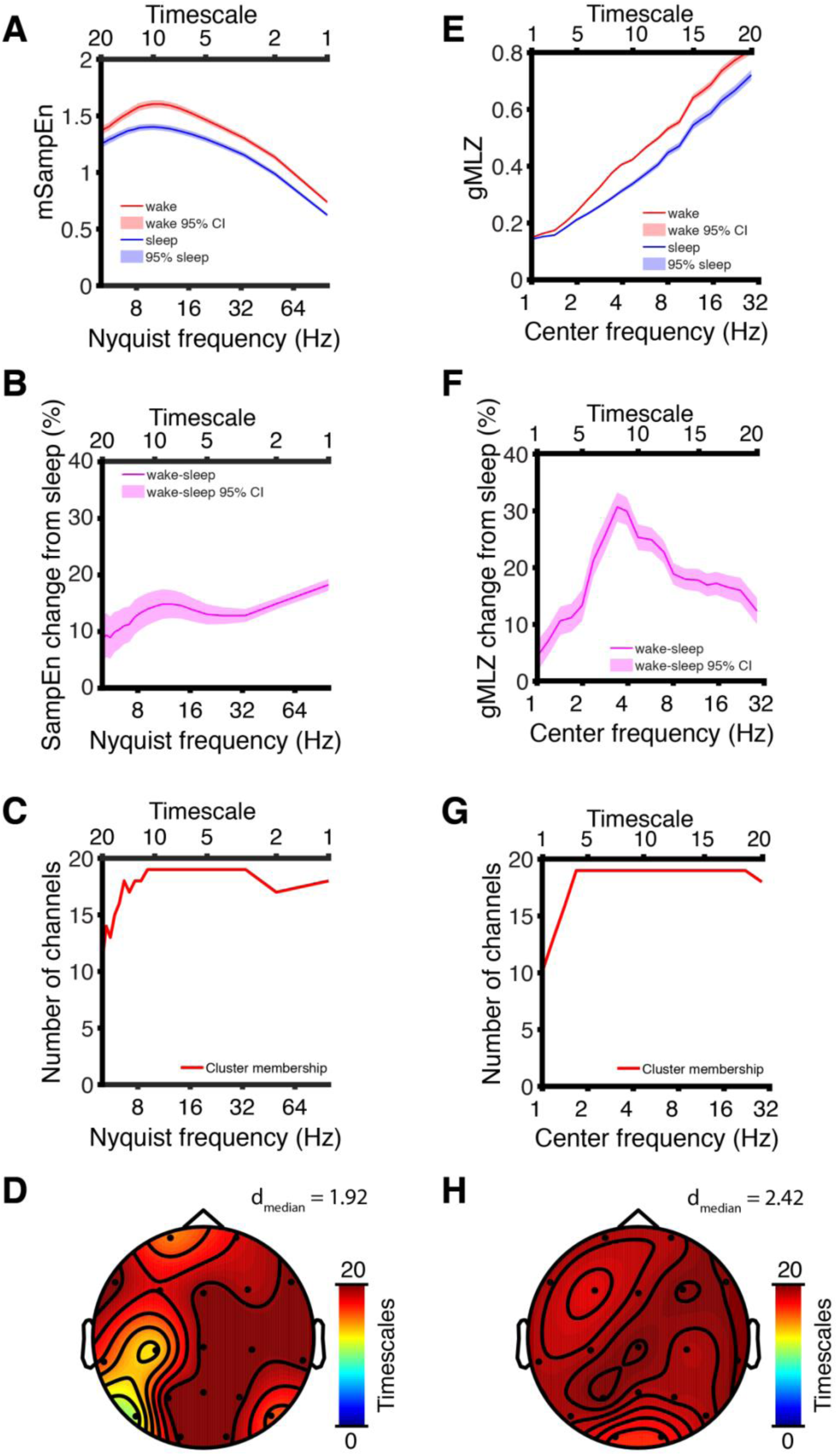
Targeted comparison of EEG complexity in sleep vs wakefulness after covarying for delta power. **(A)** Channel-averaged mMSE curves for both sleep and wakefulness show the greatest mSampEn at the 10^th^ timescale (Nyquist frequency = 10 Hz). **(B)** Percent change in channel-averaged mSampEn (wakefulness – sleep); the largest changes are observed at the 1^st^ timescales (Nyquist frequency = 100 Hz) **(C)** Membership for a channel-timescale cluster (p < 10^−4^, d_median_ = 1.92) of increased mMSE in wakefulness plotted as the number of channels participating in the cluster at each timescale. The cluster was saturated along its spatial dimension for timescales 3 – 11 (Nyquist frequency 33.3 – 9.1 Hz). **(D)** Channel-timescale cluster of increased mMSE in wakefulness color-coded by the number of timescales participating in the cluster at each channel. The cluster was nearly saturated along its timescale dimension, with its smallest extent at channels T5 and C3). **(E)** Channel-averaged gMLZ curves show greater complexity at faster timescales (i.e., timescales with shorter filter windows). **(F)** Percent change in channel-averaged gMLZ (wakefulness – sleep); complexity was greater in wakefulness, with the largest increase (30.7%) at the 8^th^ timescale corresponding to a center frequency of 3.4 Hz. **(G)** Membership for a channel-timescale cluster (p < 10^−4^, d_median_ = 2.42) of increased gMLZ in wakefulness plotted as the number of channels participating in the cluster at each timescale. The cluster was largest along its spatial dimension at timescales corresponding to delta and beta timescales (channel saturation occurred at timescales 3 – 19 corresponding to center frequencies 1.7 – 22.2 Hz). **(H)** Channel-timescale cluster of increased gMLZ in wakefulness color-coded by the number of timescales participating in the cluster at each channel. The cluster was nearly saturated along its frequency dimension.

## Discussion

Children with AS exhibit an EEG phenotype resembling states of diminished consciousness in typical individuals, while also exhibiting purposeful behavior consistent with a state of wakeful awareness, albeit marked by severe intellectual disability. This paradoxical EEG pattern during conscious wakefulness, together with similar circumstances occasionally observed in non-convulsive status epilepticus^57^, Rett syndrome^58^, and schizophrenia^59^, may be seen as a conceptual falsification of theories of consciousness which emphasize the role of information-rich brain dynamics in consciousness. At face value, the presence of pathological slow, high-amplitude, oscillations during a state of wakeful awareness is inconsistent with theoretical frameworks linking consciousness to complexity^4, 26, 27^, because diffuse hypersynchronization is informationally poor^28^ and thus lacks one of the two cardinal elements needed for a system to be conscious: information and integration^4, 26, 27^. Yet, the data presented above suggest that even in the presence of pathological, diffuse slowing of brain dynamics, the complexity of scalp EEG signals still emerges and systematically varies with levels of consciousness.

For each of three candidate biomarkers for consciousness tested (spectral power, mMSE, and gMLZ) we found significant clusters that differentiate sleep from wakefulness in AS. Significant clusters were found regardless of whether we performed a full comparison of all useable data or a targeted comparison of those EEG sections that were least likely to coincide with dream experiences (sleep state) or were especially abnormal as judged by delta EEG power (awake state). The largest within cluster effect sizes we observed (without shrinking within group variance by regressing out delta power) were those belonging to a low-frequency complexity change in the gMLZ cluster (d_median_ = 0.96/0.98 (full comparison/targeted comparison), Fig. S3A, Fig. S4A, Fig. S6A, Fig. S7A, Table S2). These effect sizes surpassed those observed for spectral power, even when considering low-frequency changes encompassed by the power cluster with the largest effects (d_median_ = 0.74/0.60 (full comparison/targeted comparison), Fig. 4C,D, Fig. S2A,B, Fig. S4B, Fig. S7C, Table 1, Table S2).

Our results clearly indicate that despite the diffuse hypersynchronized *chanting* in AS, there remains sufficient information-rich *chatter* to allow the complex dynamics typical of conscious awareness to arise. This finding resolves what would otherwise contradict views of consciousness based on informational complexity^4, 26, 27, 60^. Given these observations, how do complex brain dynamics and consciousness emerge against a background of EEG hypersynchronization? As in all scalp EEG recordings, the AS EEG is a superposition of signals from many different cortical processes and regions. To support a state of awareness, there must be a sufficient degree of complex *chatter* in AS during wakefulness, with complexity decreasing as consciousness vanishes (i.e., in sleep). Additionally, there must also be a high voltage delta *chanting* signal which drowns out the low voltage *chatter* in the AS EEG, just as one may no longer hear the *chatter* of conversation over the *chanting* of the crowd in an arena when both signals temporally coincide. To continue this analogy, the integrated energy of all *chatter* in the arena, and in the brain, may even exceed the integrated energy of the *chanting*, which, given its greater coordination, is nonetheless easier to detect^61^. The *chanting* signal in AS, however, must be functionally different from the high voltage, low-frequency activity typically observed in states of reduced consciousness^1, 6, 29^. This is because the trough of slow oscillations is believed to be associated with decreased consciousness, in sleep and anesthesia, as the system globally enters a down-state characterized by neuronal hyperpolarization^30, 31, 62^. Yet, in wakeful children with AS, consciousness does not appear to be periodically interrupted (from an observer’s perspective) as delta oscillations reach their trough. Rather, these oscillations might be more closely related to delta rhythms involved in inhibiting competing cognitive functions^63, 64^. Perhaps due to their pathologically large amplitude and diffuse nature, delta oscillations in AS might result in a broad and continuous state of cognitive inhibition, as reflected in the profound intellectual disabilities typical of this condition. For further interpretation of the AS EEG phenotype and its possible mechanisms, see Supplemental Discussion.

Although our study found that delta power was indeed modulated by diminished consciousness (i.e., sleep) in AS, the effect sizes in both comparisons (d_median_ = 0.60-0.74) were a fraction of that yielded by a prior comparison of delta power between TD control children and children with AS in the awake state (d = 1.22)^36^. Expressed as a percent change referenced to wakefulness, delta power increases when children with AS sleep by a maximum of 162% (Fig. S1B), whereas delta power in AS is greater in the awake state relative to TD control children by 1182%^36^. Thus, the difference between groups during wakefulness is an order of magnitude greater than the difference within AS with sleep/wakefulness. Because the variance within AS between conscious states is much smaller than the variance between AS and TD control children, caution should be applied when using delta EEG power alone as a biomarker for consciousness.

Finally, it is important to be mindful of some shortcomings of the present work. First, given the highly abnormal EEG presentation and the short nature of the sleep events in our data, we could not accurately perform sleep staging. Longer sessions (e.g., 24-hour recordings) might be better suited to allow an accurate sleep analysis and comparison of different stages. Furthermore, we were unable to compare sections of sleep EEG that were most and least likely to coincide with dream experiences (i.e., sleep EEG with the lowest and highest ratio of delta to high-frequency power) due to the circular nature of comparing EEG sections that are already defined such that they differ in power. Finally, while we reported a relative effect of level of consciousness on complexity metrics, a reference cohort of TD children is needed to assess the overall level of complexity present in the AS EEG.

In conclusion, this work resolves the apparent paradox of wakeful, purposefully behaving, children with AS exhibiting an EEG phenotype most typically associated with states of low/no consciousness^36, 37, 39, 65^. By finding complex brain dynamics that are sensitive to level of consciousness even under conditions of extreme cortical hypersychronization, these results support theoretical frameworks (i.e., IIT)^4, 26, 27^ linking complexity to the level of consciousness of a system. These findings, along with other rare conditions with paradoxical EEG signatures during consciousness^57–59^, warn against reverse inferring low/no consciousness in patients based on delta power^66, 67^. When brain dynamics are severely altered by genetic disorders, epilepsy, or brain injury, complexity-based methods, e.g., perturbational complexity index^9, 32, 68^, may be better suited for inferring consciousness.

## Methods

### Data acquisition

Spontaneous EEG recordings from children with AS were collected from two sites (Boston Children’s Hospital and Rady Children’s Hospital San Diego) through the AS Natural History Study [<u>NCT00296764</u>]. Consent to participate in the study was obtained from families according to the Declaration of Helsinki and was approved by the institutional review boards of the participating sites. Participants were encouraged to sleep during part of the EEG acquisition; however, due to severe sleep disturbances in AS^44^, not all children were able to fall asleep. EEG recordings were acquired in a clinical setting using an international 10-20 EEG montage (19 channels). Most participants were on central nervous system medications treating seizures or other symptoms. All EEG data were acquired at one of three native sampling rates: 250 Hz, 256 Hz, or 512 Hz. Annotations denoting sleep, drowsiness, and behavioral state were provided by the EEG technician during data acquisition. Sections of data containing drowsiness were excluded from analysis. Due to the delayed developmental abilities of many children with AS, the EEG acquisition protocol did not control for eye condition (e.g., eyes open or eyes closed) during wakefulness. Some participants gave longitudinal data across multiple visits. From a total of 161 EEG recordings from 99 participants, we identified 35 children (ages 1- 18 years) with AS whose EEG (48 recordings) contained sections of both sleep and wakefulness. Participant details are given in Table S1. Only one EEG recording was analyzed per participant according to criteria that included age and amount of good data. See Supplemental Methods for specific criteria used to select data from participants with multiple visits and for details of comparison data displayed in Fig. 1.

### Preprocessing

Data were imported to MATLAB (The MathWorks, Inc., Torrance, California) for processing and analysis. We bandpass filtered all recordings 0.5 – 45 Hz using finite impulse response filtering. Noisy channels and sections of data containing gross artifacts were manually marked to be avoided for purposes of calculating spectral power and signal complexity measures. We also omitted sections of data recorded while participants were exposed to light flash stimuli intended to trigger epileptiform activity. Stereotyped physiological and technical artifacts were removed with independent components analysis using the FastICA algorithm^69, 70^. Bad channels were spatially interpolated using a spline interpolation. A prior publication describes the full details of EEG acquisition and preprocessing^36^.

### Wavelet Transform

We computed EEG spectral power using a Morlet wavelet transform. We chose a spectral band-width of 1/2 octave (corresponding to *f/σ_f_* ∼ 8.7; *σ_f_*, spectral SD) and spaced the center frequencies logarithmically (base 2) with exponents ranging from 0 (1 Hz) to 5 (32 Hz) (inclusive) in 1/8 octave increments, yielding a total of 41 frequency bins. We then computed power in successive 3/4-overlapping temporal windows of 1 s duration. Time-frequency representations were discarded at time points where the convolution kernel overlapped with sections marked as artifact by more than 20% (see preprocessing). Finally, we averaged the time-frequency representation. Spectral power was normalized per octave, i.e., log_2_(Hz), rather than Hz to account for the logarithmic nature of EEG signals^71^. To plot spectral power, we first averaged across channels, then log-transformed power before averaging across participants. For statistical comparisons between groups, we log-transformed power at each point in channel-frequency space (see Statistical Analysis below).

### Signal Complexity

We measured EEG signal complexity using two methods, mMSE^48, 49^ and gMLZ^50–52^ complexity. For further details on how each measure is computed, see the Supplemental Methods. To compute mMSE, we used 30 s segments with 20 coarse graining scales, thus allowing for as many as 300 samples at the 20^th^ timescale. This number of timescales gives good coverage of the EEG spectrum, with a Nyquist frequency of 100 Hz for the 1st timescale and a Nyquist frequency of 5 Hz for the 20^th^ timescale. Following the advice of Grandy and colleagues^72^, we rejected all segments that did not include at least 100 valid samples for each timescale. To compute gMLZ, we used smaller segments for reasons of computational feasibility. Ibáñez-Molina and colleagues^51^ found no benefit to using EEG segments longer than 2000 samples for computing MLZ. For this reason, we used 12 s (2400 sample) EEG segments for computing gMLZ, exceeding the recommendation of Ibáñez-Molina and colleagues to afford elbow room for EEG segments with excised artifacts. The gMLZ derived from each EEG segment was normalized according to the number of valid samples n using the quantity n/log_2_(n)^50^. For each timescale, two moving median filters were employed, one to smooth the EEG signal itself and another to compute the dynamic threshold that is used to binarize the signal^52^. We utilized 20 timescales with logarithmically spaced center frequencies 1 – 30 Hz. See Table 2 for gMLZ center frequencies, smoothing window sizes, and bandwidths at each timescale.

### Comparison of Sleep Versus Wakefulness

Our comparison of data from sleep and wakefulness is informed by the finding that most awakenings from NREM sleep are accompanied by reports of dreams^56^ (and are thus “contaminated” by consciousness). In addition to the variance in level of consciousness encountered in sleep, there is large variance in delta amplitude encountered during wakefulness in AS^37^. For these two reasons, we performed two comparisons of EEG data: 1) a full comparison using all good data from both the awake and the asleep state and 2) a targeted comparison using sections of sleep EEG that are unlikely to coincide with conscious experience (as judged by parietal EEG activity) paired with sections of awake EEG that are especially abnormal as judged by their delta power. For a detailed explanation of how data sections were chosen for the targeted comparison, see Supplemental Methods.

### Statistical Analysis

Both mMSE and gMLZ are defined such that regularities (e.g., delta oscillations) in the signal diminish complexity. For this reason, we covaried for delta (1 – 4 Hz integrated) power using simple linear regression models (separate model for each channel, timescale, and complexity measure). We report results of the targeted comparison both with and without covarying for delta power. To infer whether changes in signal complexity between sleep and wakefulness were mediated by changes in delta power, we used a nonparametric (2 x 10^4^ bootstraps) path analytic framework for mediation analysis^73^.

To account for a large number of comparisons across channels and timescales or frequencies, we used permutation cluster statistics to test for differences between sleep and wakefulness in both complexity and spectral power^53, 54^. We first performed t-tests at each channel and scale/frequency and then thresholded t-statistics using p = 0.01 before clustering in channel-scale space (complexity) or channel-frequency space (spectral power). For each cluster, we then derived two-tailed statistical significance nonparametrically by permuting the condition labels 10^4^ times and comparing the size of the original cluster to the empirical distribution of cluster sizes. This approach is unbiased with respect to directionality, frequency/timescale, and electrode location. In total, we performed 8 separate tests: power and complexity were examined in both a full comparison and a targeted comparison (3 EEG measures x 2 comparisons), and complexity was also examined in a follow-up targeted comparison in which we covaried for delta power (2 EEG measures x 1 comparison). We then adjusted for the total number of tests using a Bonferroni correction, yielding α = 0.0063. Effect sizes for each cluster (median across all cluster members) were measured as Cohen’s d.

## Grants and acknowledgements

This work was made possible by the US National Institutes of Health (NIH) funded AS Natural History Study [<u>NCT00296764</u>], NIH Grant Nos. <u>U54RR019478</u> (LMB; principal investigator, Arthur Beaudet) and <u>U54HD061222</u> (LMB; principal investigator, Alan Percy), and funding from the Tiny Blue Dot (TBD) foundation (MMM). We sincerely thank all children and families who participated in this work. We also thank Melanie Boly and Anthony Hudetz for providing code for the Lempel-Ziv algorithm. Finally, we thank Shafali S. Jeste for contributing TD EEG in Fig. 1B and Paul Vespa for contributing vegetative state EEG in Fig. 1G.

## Contributions

JF and MMM conceived the concept. LMB collected the data and provided clinical expertise. JFH created the EEG processing pipeline tools. JFH and JF both wrote code for the analysis. JF processed and analyzed the data. JD provided computing resources. MAJ performed the mediation analysis. JF and MMM wrote the manuscript. All authors discussed the results and interpretation and commented on the manuscript.

## Supplemental material

### Supplemental Methods

#### EEG data selection

In cases where participants gave data at multiple visits, we analyzed EEG from the visit that yielded the greatest number of valid frequency transform windows at the lowest frequency analyzed (1 Hz) in data sections from the targeted comparison. Ties were broken using the youngest visit, as delta power is known to attenuate with age in AS^1^. Because the sampling rate influences multiscale analyses, all EEG signals were downsampled to 200 Hz without filtering prior to computing mMSE and gMLZ. Data sections containing artifacts, drowsiness, or light flashes were excised prior to computing mMSE and gMLZ. In the full comparison, mMSE and gMLZ were computed in nonoverlapping segments; in the targeted comparison, we applied 50% overlap between data segments to give better coverage of shorter data.

Our comparison of data from sleep and wakefulness is informed by the finding that most awakening from NREM sleep are accompanied by reports of dreams^2^ (and are thus “contaminated” by consciousness). Siclari and colleagues recently found that dreams are most likely to be reported following sleep in which delta power is low and high-frequency (20 – 50 Hz) power is high over a posterior hot zone (PHZ)^3^. Conversely, the same study found that sections of sleep characterized by high delta power and low high-frequency power in the PHZ are likely to coincide with no reportable conscious experience. Moreover, the PHZ encompasses the precuneus, a parietal area heavily implicated in conscious awareness^4–6^. The ability to stratify sleep according to sections that are more likely and less likely to correspond to conscious experience motivates a more targeted comparison of asleep and awake EEG in AS.

In addition to the variance in level of consciousness encountered in sleep, there is large variance in delta amplitude encountered during wakefulness in AS. The awake state AS delta EEG phenotype shows greater dynamic variability (i.e., intermittent bursts of delta activity) across all scalp regions as compared with TD control children^7^. For these reasons, we performed two comparisons of EEG data: 1) a full comparison using all good data from both the awake and the asleep state and 2) a targeted comparison using sections of sleep EEG that are unlikely to coincide with conscious experience (as judged by parietal EEG activity) paired with sections of awake EEG that are especially abnormal as judged by their delta power.

Data sections for the targeted comparison were identified separately for awake and asleep state data by partitioning the time series (awake: delta power; asleep: parietal delta/fast power) into regions that minimize the sum of square difference between each sample and its local mean. In each case, the maximum number of breaks between regions was not allowed to exceed the length of usable EEG data in minutes for the given condition. For sleep data, we optimized the ratio of delta power (integrated 1-4 Hz) over high-frequency power (integrated 20 – 45 Hz) averaged across parietal channels (Pz, P3, and P4) from the time-frequency representation of spectral power. For awake data, we optimized delta power (integrated 1-4 Hz) averaged across all channels from the time-frequency representation of spectral power. In each condition, we started with the region with the highest mean and continued selecting additional regions with the next highest means until the combined length of all regions met or exceeded a fixed amount determined as a function of the total data length. These fixed amounts were as follows: 30 s (for total data length ≤ 60 s), 60 s (total length ≤ 120 s), 90 s (≤ 300 s), 120 s (total length ≤ 600 s), 180 s (total length ≤ 1200 s), 240 s (total length > 1200 s). EEG data from these regions were then entered into the targeted comparison.

#### Signal complexity

The original multiscale entropy (MSE) was introduced by Costa and collogues^8^ using sample entropy (SampEn)^9^, or the tendency for short motifs to reoccur in a signal within a tolerance r (defined as a fixed proportion of the signal’s standard deviation). Here we used mMSE (r = 0.15), an improved version of the original MSE algorithm based on the more robust mSampEn^10^, which is less sensitive to r. We implemented mMSE using custom code that patches a commonly cited shortcoming of the original MSE algorithm^11, 12^ by computing r separately for each timescale.

As an additional complexity measure, we also examined the number of substrings contained in the binarized EEG signal^13^ using gMLZ. A multiscale approach to Lempel-Ziv complexity was first advocated by Ibáñez-Molina and colleagues^14^, who showed that a dynamic threshold applied with different smoothing windows for different timescales shows better sensitivity to all EEG frequencies than a static threshold obtained from the median of the entire signal, which is biased towards lower EEG frequencies. More recently, however, Yeh and Shi^15^ demonstrated that the thresholding approach advocated by Ibáñez-Molina and colleagues may in fact be too biased toward higher EEG frequencies, thus risking overestimates of complexity. As a solution, Yeh and Shi have proposed gMLZ, which uses the same dynamic threshold as the approach given by Ibáñez-Molina and colleagues while also applying a moving median filter with a smaller smoothing window to the signal before binarizing the signal according to its threshold given by a moving median filter with a larger smoothing window. We implemented gMLZ by modifying existing code provided by Hudetz and colleagues that implements the Lempel-Ziv algorithm^16^.

#### Comparison data

For illustrative purposes, we qualitatively compared AS EEG waveforms with EEG waveforms from external data corresponding to conscious and unconscious states (Fig. 1). AS EEG waveforms were displayed from participant 10 of this study (Table S1), with representative sections extracted and display from wakefulness (Fig. 1A) and sleep (Fig. 1E). This participant was not on any medications and did not have epilepsy. Awake state EEG data (referential recording) from a typically developing 38-month-old girl were provided courtesy of Shafali Jeste at the University of California, Los Angeles^17^. The girl’s EEG data were acquired as control data for research purposes with her family’s consent. A representative section of this girl’s EEG was displayed for comparison with AS (Fig. 1B). EEG data (bipolar recordings) from a 37-year-old woman without sleeping problems^18^ (Fig. 1C,D,H) and a 2-year-old girl with epilepsy^19^ (Fig. 1F) were downloaded from PhysioNet, an National Institutes of Health funded collection of publicly available physiological signals^20^. Annotations provided by PhysioNet were used to extract and display representative sections of EEG recorded during wakefulness (Fig. 1C), REM sleep (Fig. D), and stage 4 NREM sleep (Fig. 1H) from the 37-year-old woman, and of ictal EEG recorded during an epileptic seizure (Fig. 1 F) from the 2-year-old girl. Clinical EEG data from a 59-year-old man in a vegetative state were provided courtesy of Paul Vespa at the University of California, Los Angeles. This patient was admitted to an intensive care unit after sustaining brain injury during a fall down the stairs (Glasgow Coma Scale = 3, post-resuscitation) and was later discharged from the intensive care unit in a vegetative state (Extended Glasgow Outcome Scale = 2). The patient was consented by his family to participate in research. A representative portion of this patient’s EEG data was extracted and displayed (Fig. 1G).

### Supplemental discussion

Children with AS display hypersynchronous delta activity suggestive of hyperpolarized cortical down states while nonetheless exhibiting wakeful, conscious behavior. While such activity is observed at all scalp electrodes, it is untenable to assume that the entire AS cortex is hypersynchronized while children are awake and behaving. An alternative explanation for the AS EEG phenotype is that such hypersynchronous activity could be driven by a focal source whose activity conducts broadly to all scalp regions. In this scenario, hypersynchronization is local, sparing most of cortex, but appears global due to volume conduction. A similar scenario may occur in other conditions, such as disorders of consciousness resulting from TBI, in which injured cortical tissue becomes pathologically synchronous at a focal site^21–24^, plausibly because of reactive astrocytes that exacerbate neuronal excitability^25, 26^. Nonetheless, the globally diffuse topography of wakeful delta power in AS does not obviously suggest a focal source^1^, and global delta coherence in AS has recently been shown to be similar to that in TD children during both wakefulness and sleep^27^. The foregoing suggests that, while theoretically possible, the single focal generator hypothesis is also implausible.

To further elucidate the mechanism and anatomical substrates of the AS EEG phenotype, EEG source localization estimates should be obtained from children with AS. Future work should also examine intracranial electrophysiology in animal models of AS (e.g., *Ube3a* knock-out mice) to assess the extent of global hypersynchronization in AS cortex. Local field potential (LFP) recordings from a mouse model of AS have shown hypersynchronous delta activity in Layer 4 of V1 while mice are awake, head-fixed, and unanaesthetized^7, 28^. However, as argued above, a focal source in V1 alone is unlikely to explain the diffuse hypersynchronous delta phenotype observed in the AS scalp EEG.

## Supplemental Figures

**Figure S1.**
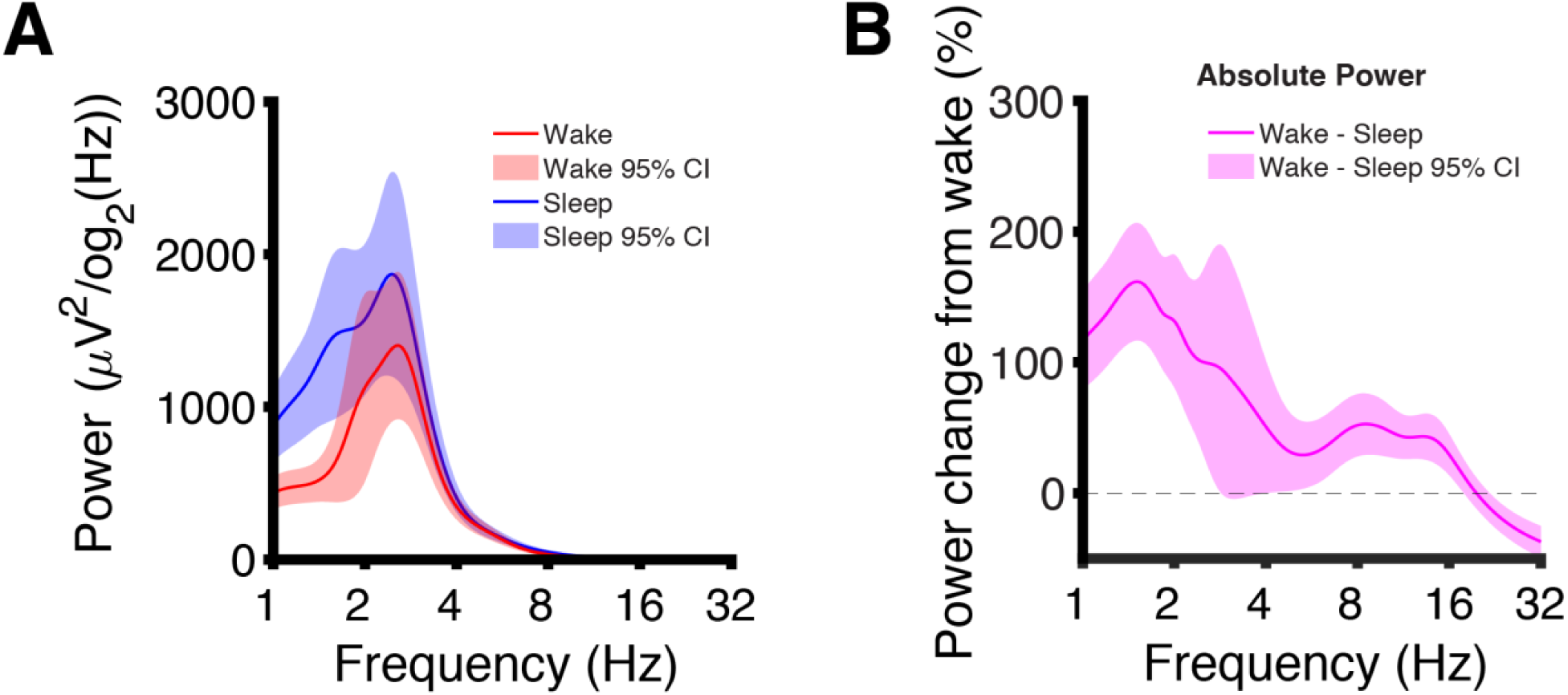
Alternative visualizations of EEG change from the full comparison **(A)** Channel-averaged untransformed EEG power traces demonstrate global maxima in the delta band for both sleep and wakefulness. This untransformed view of the data (without log-scaling) allows for a clearer visualization of the delta band. The delta EEG power in sleep shows a peak which is broader on the left side (with more power in the 1.0 – 1.6 Hz band than in wakefulness as judged by nonoverlapping 95% confidence intervals). Although the delta peak in the asleep state is broad, it features only one local maximum (cf. Fig. S6A). **(B)** Channel-averaged EEG power change referenced to wakefulness. The largest power increase in sleep occurs at f = 1.51 Hz (162% increase), and the largest reduction in sleep occurs at f = 32 Hz (37.1% decrease).

**Figure S2.**
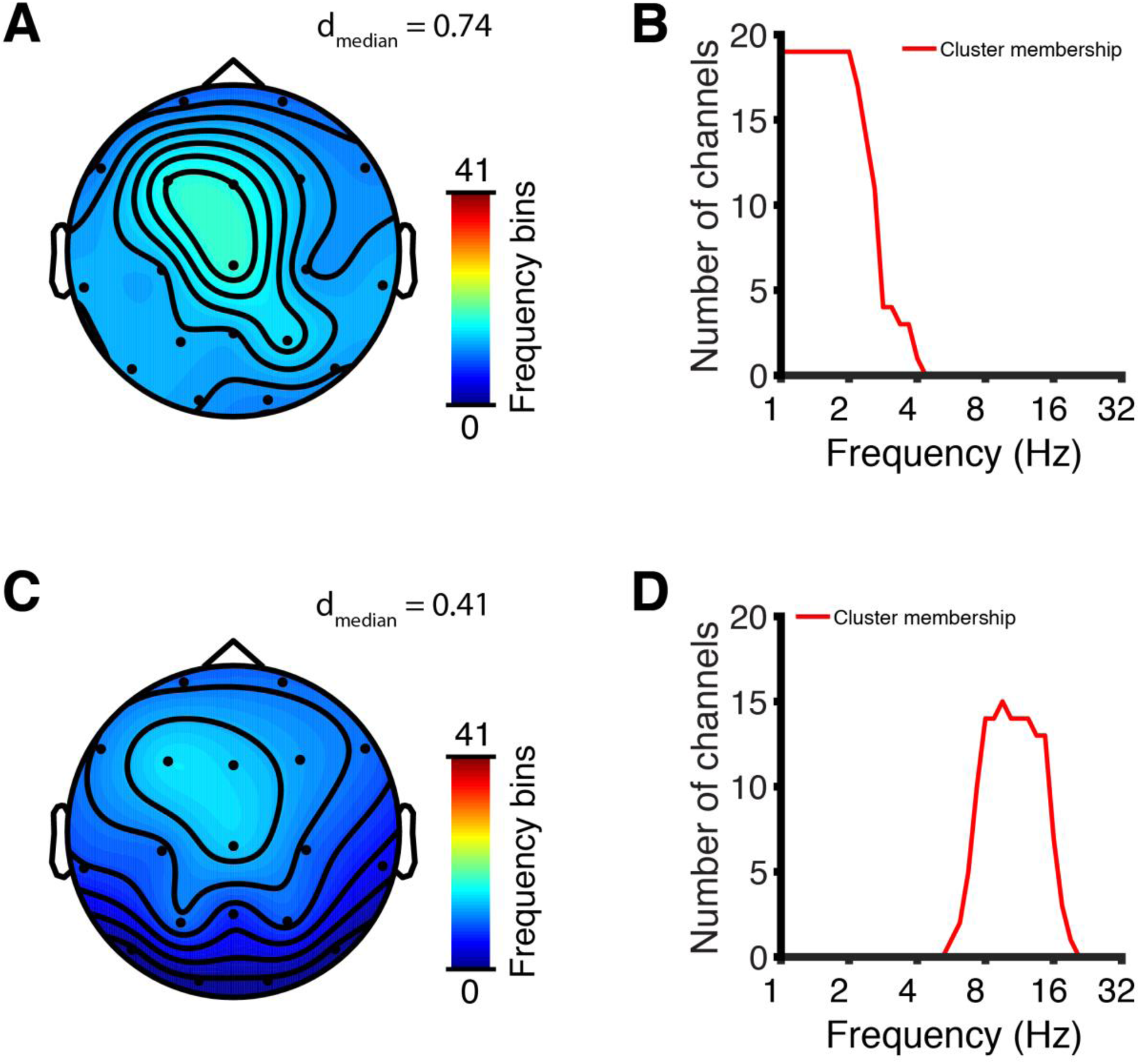
Channel-frequency clusters of decreased power in wakefulness derived from permutation cluster statistics using a stricter threshold (p = 0.0005) for clustering t-statistics as a means of breaking apart a cluster with two oscillatory changes that were fused at a more permissive threshold (p = 0.01, Fig. 2C,D) in the full comparison. **(A)** Channel-frequency cluster (p < 10^−4^, d_median_ = 0.74) of decreased power in wakefulness color-coded by the number of frequency bins participating in the cluster at each channel. The cluster was largest along its frequency dimension over frontocentral scalp regions. **(B)** Channel-frequency cluster membership plotted as the number of channels participating in the cluster at each frequency bin. The cluster was saturated along its spatial extent at frequencies ≤ 1.0 Hz. **(C)** Channel-frequency cluster (p < 10^−4^, d_median_ = 0.41) of decreased power in wakefulness color-coded by the number of frequency bins participating in the cluster at each channel. The cluster was largest along its frequency dimension over frontocentral scalp regions. **(D)** Channel-frequency cluster membership plotted as the number of channels participating in the cluster at each frequency bin. The cluster was largest along its spatial extent at alpha and low-beta frequencies (8 – 14 Hz).

**Figure S3.**
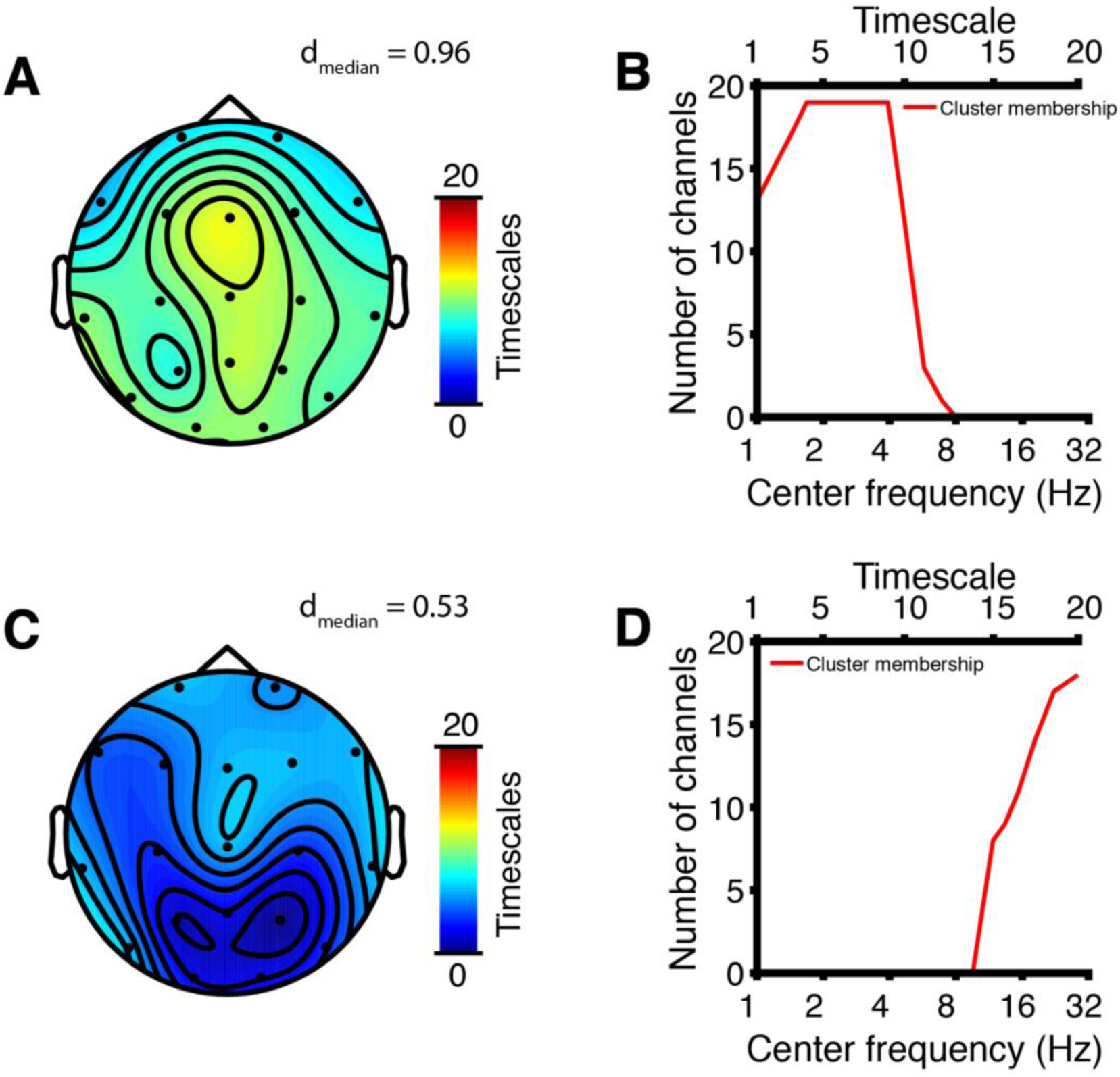
Channel-timescale clusters of increased gMLZ in wakefulness derived from permutation cluster statistics using a stricter threshold (p = 0.0005) for clustering t-statistics as a means of breaking apart a cluster with two complexity changes that were fused at a more permissive threshold (p = 0.01, Fig. 3G,H) in the full comparison. **(A)** Channel-timescale cluster (p < 10^−4^, d_median_ = 0.96) of increased gMLZ in wakefulness color-coded by the number of timescales participating in the cluster at each channel. The cluster was largest along its timescale dimension over channel Fz and, like the mMSE cluster identified in Fig. 3D, encompassed the fewest timescales at the most anterior channels (F7, Fp1, Fp2, and F8). **(B)** Channel-timescale cluster membership plotted as the number of channels participating in the cluster at each timescale. The cluster was saturated along its spatial extent at delta frequencies (f = 1.68 – 3.92 Hz). **(C)** Channel-timescale cluster (p = 0.0004, d_median_ = 0.53) of increased gMLZ in wakefulness color-coded by the number of timescales participating in the cluster at each channel. The cluster exhibited local minima (fewest timescales) over channels P3 and P4 (cf. Fig. 3H). **(D)** Channel-timescale cluster membership plotted as the number of channels participating in the cluster at each timescale. The cluster largely encompassed beta frequencies (f = 11.8 – 28.6 Hz).

**Figure S4.**
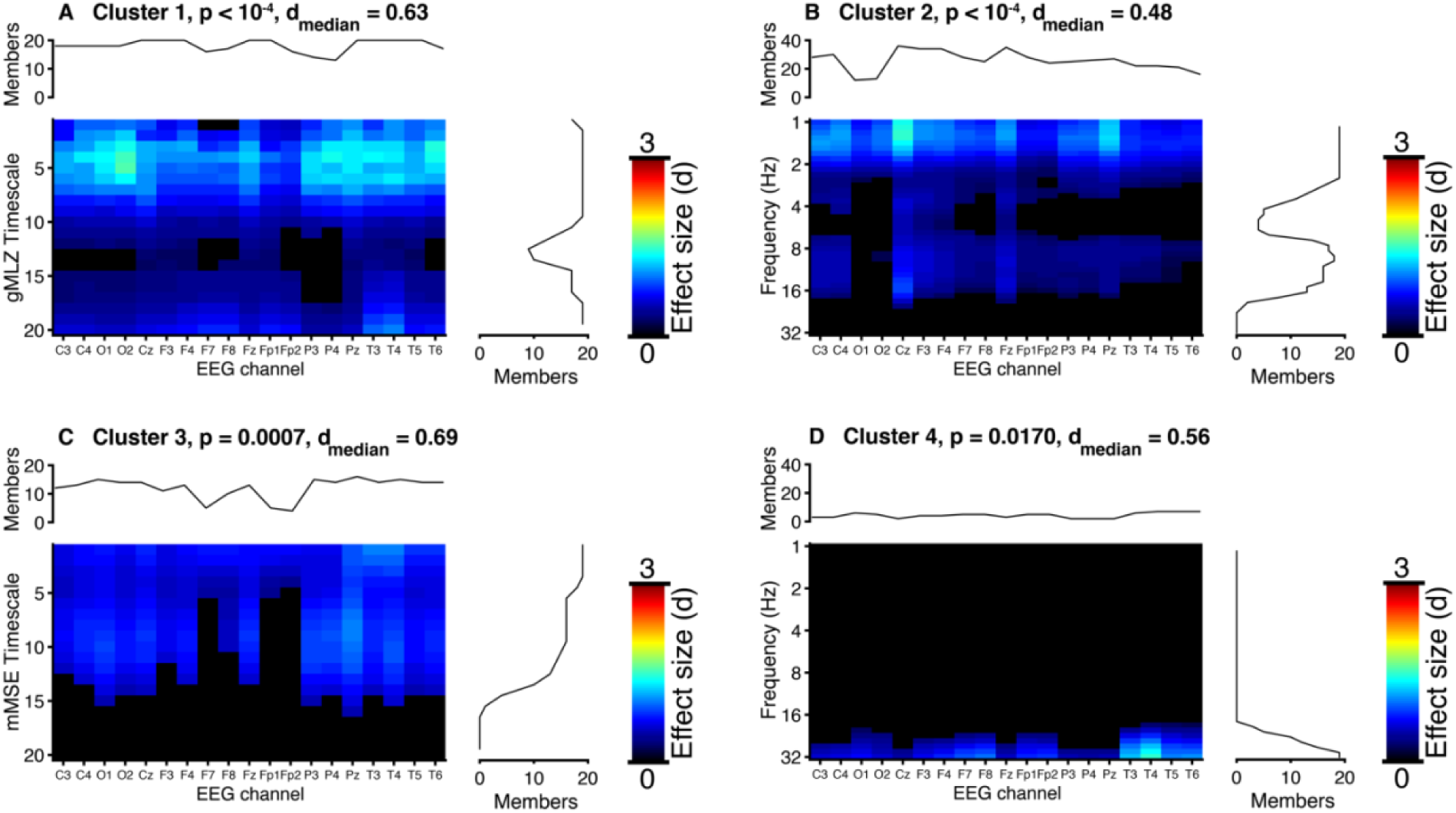
Channel-timescale/frequency space clusters from the full comparison. Heatmaps reflect the unsigned effect size (Cohen’s d); flanking graphs count cluster membership. Cluster numbers refer to Table 1. Permutation cluster statistics correct for multiple testing across channels and timescales/frequencies, while a Bonferroni correction is used to correct for multiple testing across analyses and EEG measures. Statistical significance is determined using α = 0.0063 (Bonferroni correction). **(A)** Significant gMLZ cluster (greater in wakefulness) covering 90.79% of channel-timescale space (see Fig. 3G,H). **(B)** Significant power cluster (greater in sleep) covering 62.39% of channel-frequency space (see Fig. 2C,D). **(C)** Significant mMSE cluster (greater in wakefulness) covering 60.79% of channel-timescale space (see Fig. 3C,D). **(D)** Power cluster (greater in wakefulness) covering 10.65% of channel-frequency space (not significant after Bonferroni correction).

**Figure S5.**
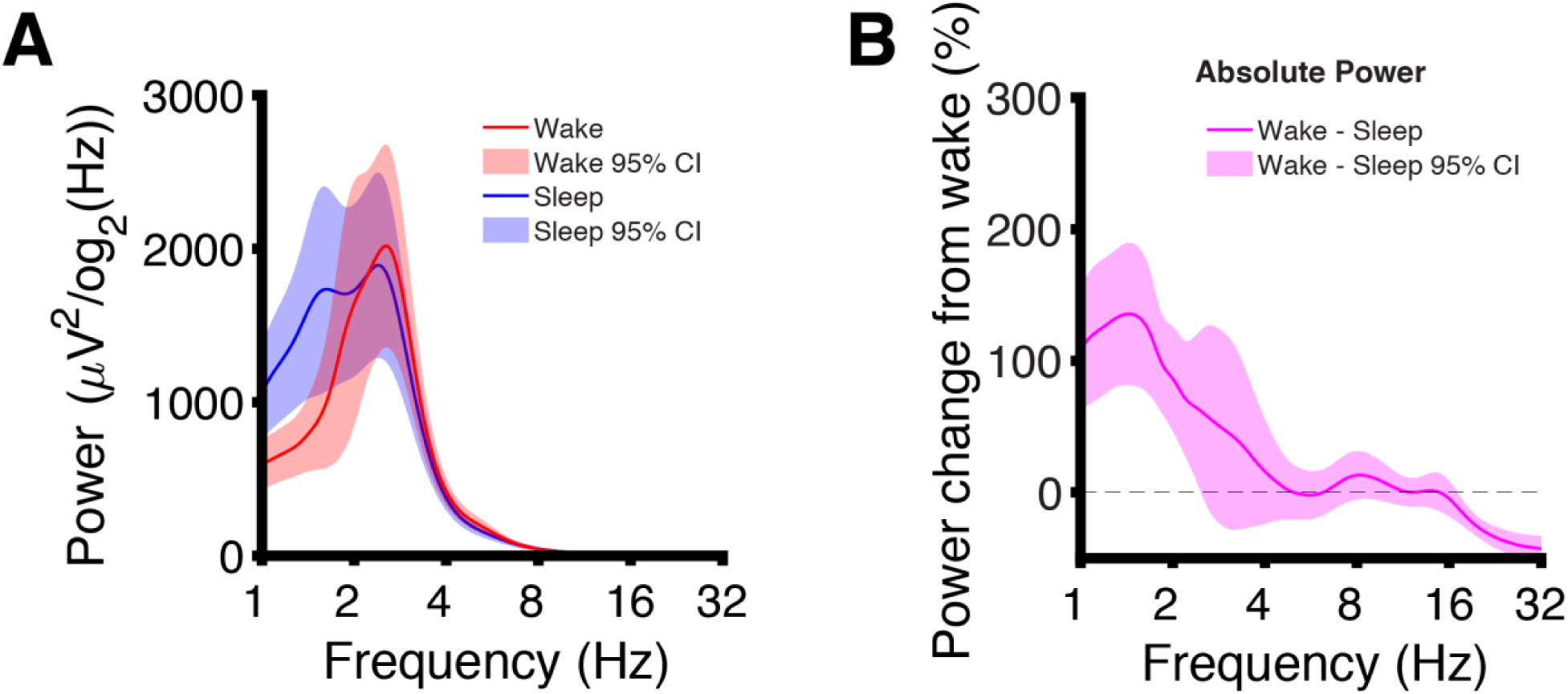
Alternative visualizations of EEG power from the targeted comparison **(A)** Channel-averaged untransformed EEG power traces (targeted comparison) demonstrate global maxima in the delta band for both sleep and wakefulness. This untransformed view of the data (without log-scaling) allows for a clearer visualization of the delta band. The EEG power in sleep shows two peaks in the delta band (f_1_ = 1.62 Hz, f_2_ = 2.40 Hz), whereas only one delta peak is present in wakefulness (f = 2.55 Hz). This suggests two separate oscillatory processes for slow waves in sleep and for the AS EEG phenotype. Note that because the log-transform is a nonlinear transform, peak frequencies differ between untransformed and log-scaled power (cf. Fig. 4A). The asleep EEG exhibits more power in the 1.0 – 1.4 Hz band than the awake EEG as judged by 95% confidence intervals. **(B)** Channel-averaged EEG power change referenced to wakefulness. The largest power increase in sleep occurs at f = 1.44 Hz (136% increase), and the largest reduction in sleep occurs at f = 32 Hz (42.7% decrease).

**Figure S6.**
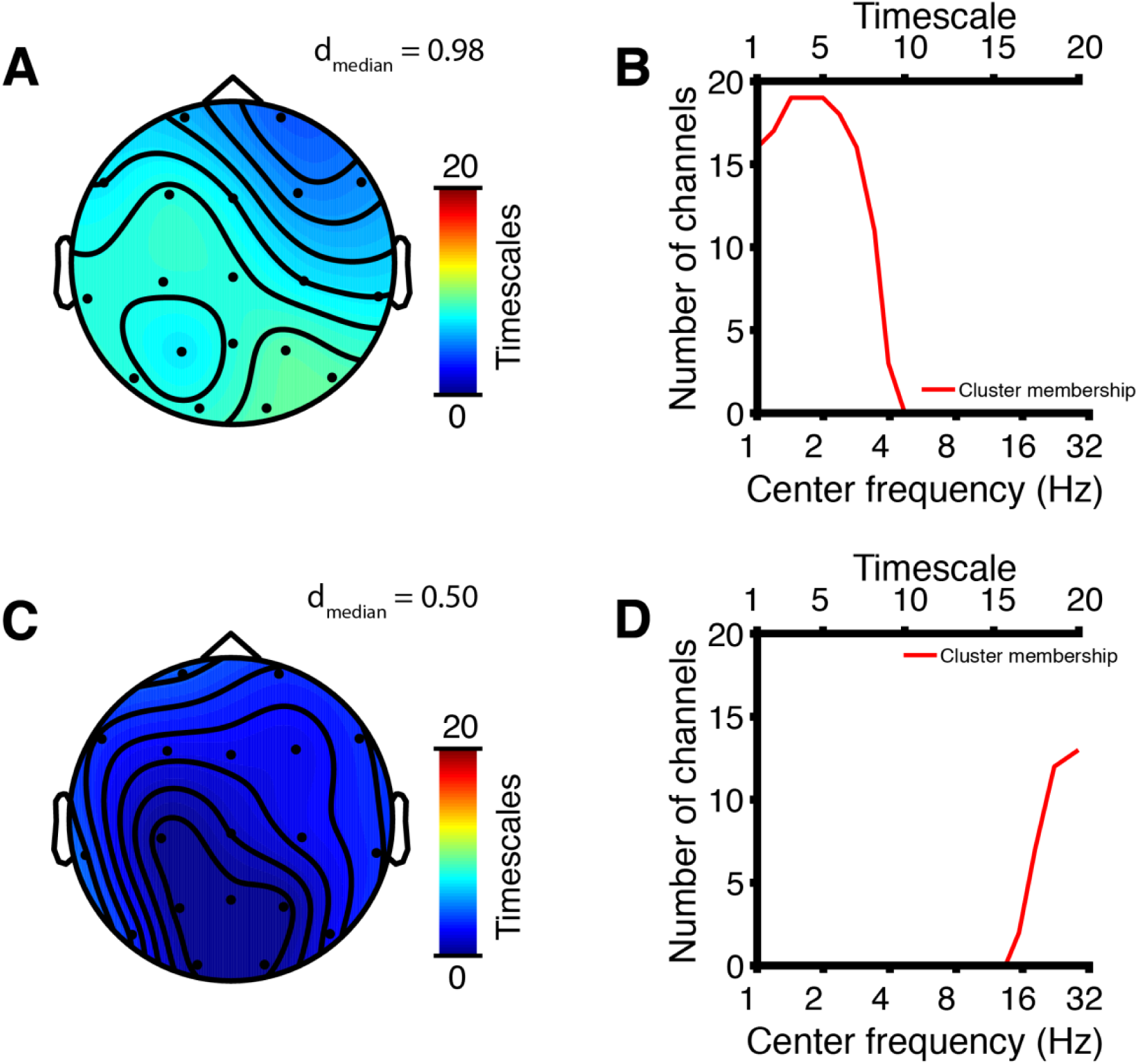
Channel-timescale clusters of increased gMLZ in wakefulness derived from permutation cluster statistics using a stricter threshold (p = 0.0005) for clustering t-statistics as a means of breaking apart a cluster with two complexity changes that were fused at a more permissive threshold (p = 0.01, Fig. 5G,H) in the targeted comparison. **(A)** Channel-timescale cluster (p = 0.0001, d_median_ = 0.98) of increased gMLZ in wakefulness color-coded by the number of timescales participating in the cluster at each channel. The cluster was smallest along its timescale dimension over right frontal scalp regions. **(B)** Channel-timescale cluster membership plotted as the number of channels participating in the cluster at each timescale. The cluster was saturated along its spatial extent at f = 1.42 – 1.98 Hz. **(C)** Channel-timescale cluster (p = 0.0013, d_median_ = 0.50) of increased gMLZ in wakefulness color-coded by the number of timescales participating in the cluster at each channel. The cluster was smallest along its timescale dimension over posterior scalp regions. **(D)** Channel-timescale cluster membership plotted as the number of channels participating in the cluster at each timescale. The cluster largely encompassed beta frequencies (f = 15.4 – 28.6 Hz).

**Figure S7.**
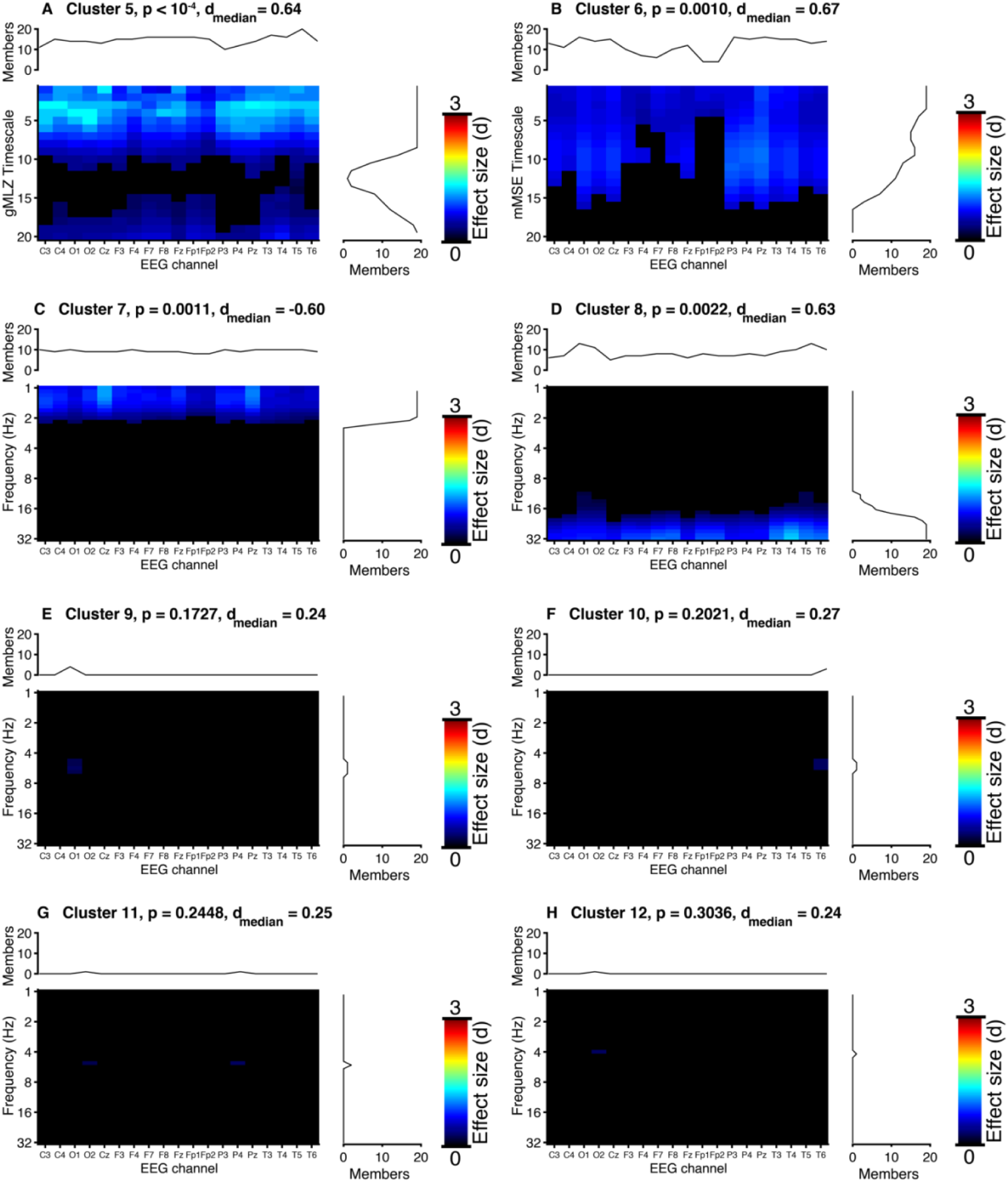
Channel-timescale/frequency-channel space clusters from the targeted comparison. Heatmaps reflect the unsigned effect size (Cohen’s d); flanking graphs count cluster membership. Cluster numbers refer to Table 1. Permutation cluster statistics correct for multiple testing across channels and timescales/frequencies, while a Bonferroni correction is used to correct for multiple testing across analyses and EEG measures. Statistical significance is determined using α = 0.0063 (Bonferroni correction). **(A)** Significant gMLZ cluster (greater in wakefulness) covering 73.42% of channel-timescale space (see Fig. 5G,H). **(B)** Significant mMSE cluster (greater in wakefulness) covering 59.47% of channel-timescale space (see Fig. 5C,D). **(C)** Significant power cluster (greater in sleep) covering 22.72% of channel-frequency space (see Fig. 4C,D). **(D)** Significant power cluster (greater in wakefulness) covering 20.15% of channel-frequency space (see Fig. 4E,F). **(E)** Power cluster (greater in wakefulness, not significant) covering 0.51% of channel-frequency space. **(F)** Power cluster (greater in wakefulness, not significant) covering 0.39% of channel-frequency space. **(G)** Power cluster (greater in wakefulness, not significant) covering 0.26% of channel-frequency space. **(H)** Power cluster consisting of only 1 point (greater in wakefulness, not significant) covering 0.13% of channel-frequency space.

**Figure S8.**
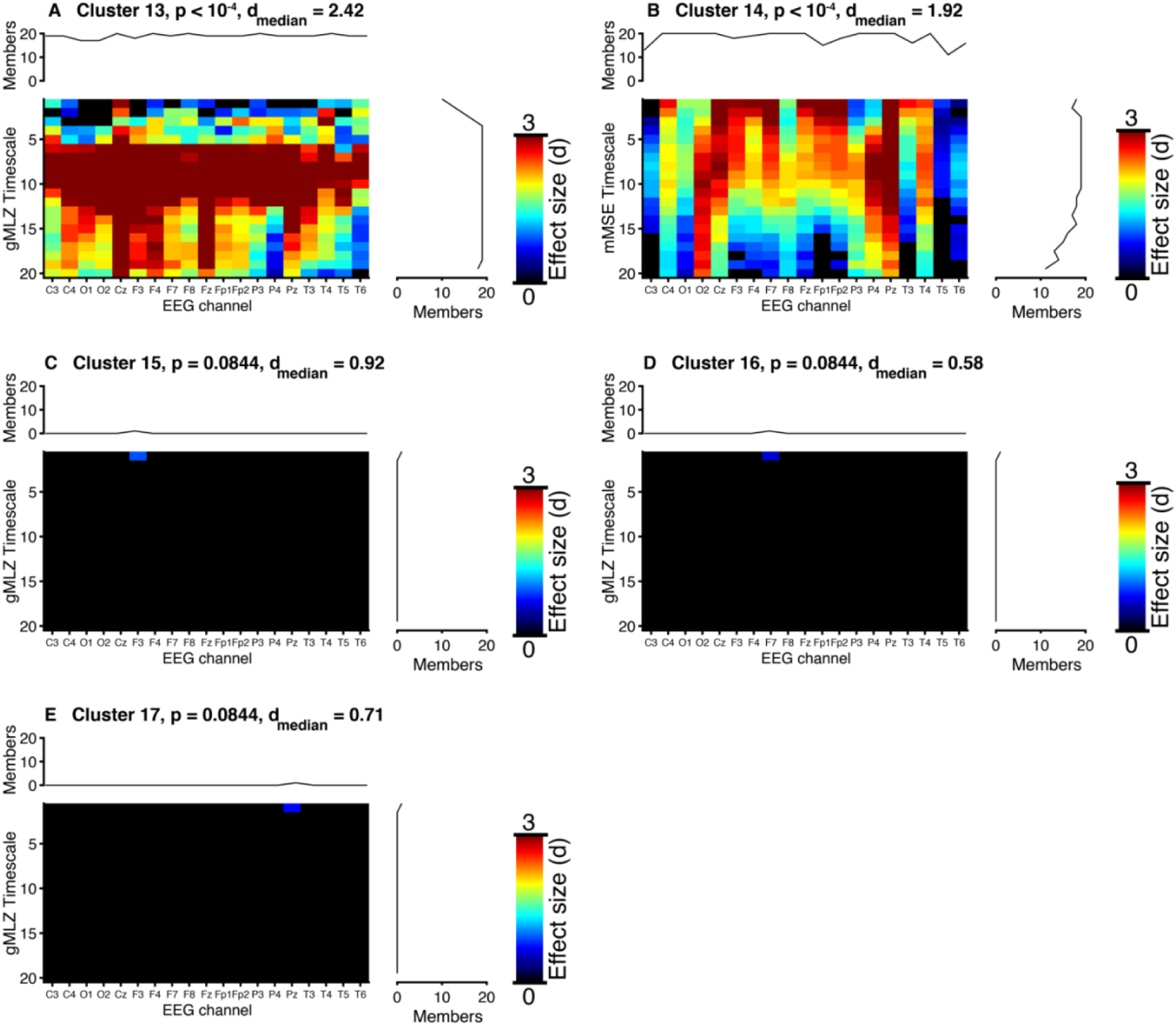
Timescale-channel space clusters from the targeted comparison covarying for delta power. Heatmaps reflect the unsigned effect size (Cohen’s d); flanking graphs count cluster membership. Cluster numbers refer to Table 1. Permutation cluster statistics correct for multiple testing across channels and timescales/frequencies, while a Bonferroni correction is used to correct for multiple testing across analyses and EEG measures. Statistical significance is determined using α = 0.0063 (Bonferroni correction). **(A)** Significant gMLZ cluster (greater in wakefulness) covering 95.00% of channel-timescale space (see Fig. 6I,J). **(B)** Significant mMSE cluster (greater in wakefulness) covering 91.05% of channel-timescale space (see Fig. 6D,E) **(C)** gMLZ cluster consisting of only 1 point (greater in sleep, not significant) covering 0.26% of channel-timescale space. **(D)** gMLZ cluster consisting of only 1 point (greater in sleep, not significant) covering 0.26% of channel-timescale space. **(E)** gMLZ cluster consisting of only 1 point (greater in sleep, not significant) covering 0.26% of channel-timescale space.

## Supplemental Tables

**Table S1.** AS cohort demographics and data. Participants are sorted by age and described according to sex, genotype (i.e., the presence of absence of a 15q deletion), seizure status, and medication. The amount of data included in both the full analysis and targeted analysis is also given for each participant. For the targeted analysis, data length is described both in terms of data length (in seconds) and number of continuous data segments. 27 out of 35 (77%) participants were on at least one medication acting on the central nervous system (CNS, includes over-the-counter medications such as melatonin). 27 participants (overlapping with but not identical to the subset on medications) also had a history of seizures at the time of EEG recording.

|  | Age<br>(months) | Sex | 15q11.2-q13.1<br>deletion? | Seizures? | CNS<br>meds? | Awake data, full<br>comparison (s) | Awake data,<br>targeted<br>comparison (s) | Awake<br>sections,<br>targeted<br>comparison | Asleep data,<br>full<br>comparison | Asleep data,<br>targeted<br>comparison<br>(s) | Asleep sections,<br>targeted<br>comparison |
| --- | --- | --- | --- | --- | --- | --- | --- | --- | --- | --- | --- |
| Participant 1 | 13 | female | Yes | No | No | 1561.5 | 566.6 | 7 | 461.8 | 145.2 | 5 |
| Participant 2 | 13 | female | No | Yes | Yes | 10019 | 438.1 | 5 | 7362.1 | 253.4 | 5 |
| Participant 3 | 15 | male | Yes | Yes | No | 372 | 163.5 | 2 | 1635.5 | 266.3 | 9 |
| Participant 4 | 16 | female | No | Yes | Yes | 664.5 | 193 | 5 | 977.2 | 305.3 | 7 |
| Participant 5 | 16 | male | Yes | No | No | 470.7 | 149 | 2 | 989.2 | 182.1 | 5 |
| Participant 6 | 22 | male | Yes | No | Yes | 264.2 | 192.3 | 4 | 1695.1 | 246.1 | 7 |
| Participant 7 | 24 | male | Yes | Yes | Yes | 615.8 | 221.6 | 3 | 1804.9 | 288.1 | 7 |
| Participant 8 | 25 | male | Yes | Yes | Yes | 238.5 | 144.5 | 2 | 1585.3 | 261.9 | 9 |
| Participant 9 | 26 | female | Yes | Yes | Yes | 822.3 | 237.5 | 5 | 319.6 | 148.3 | 2 |
| Participant 10 | 27 | female | Yes | No | No | 593 | 207 | 2 | 454.7 | 153.6 | 3 |
| Participant 11 | 29 | male | No | No | Yes | 367.6 | 202 | 2 | 642.2 | 143 | 5 |
| Participant 12 | 33 | female | Yes | Yes | Yes | 227.1 | 126.8 | 1 | 630.4 | 142.6 | 3 |
| Participant 13 | 34 | male | Yes | Yes | Yes | 720.8 | 195 | 2 | 1338.4 | 257.5 | 7 |
| Participant 14 | 36 | female | Yes | No | Yes | 1259 | 350.7 | 4 | 665.3 | 184.6 | 3 |
| Participant 15 | 37 | male | Yes | Yes | No | 513.1 | 172.8 | 3 | 1774.4 | 241.1 | 7 |
| Participant 16 | 46 | female | No | No | Yes | 339.6 | 124.4 | 3 | 227.1 | 96.6 | 2 |
| Participant 17 | 47 | male | No | Yes | Yes | 431.4 | 195.2 | 3 | 1251.1 | 181.1 | 7 |
| Participant 18 | 47 | male | Yes | Yes | Yes | 1026.1 | 180.9 | 3 | 709.6 | 196.9 | 5 |
| Participant 19 | 49 | male | No | Yes | Yes | 1048.5 | 208.5 | 5 | 769.9 | 245 | 3 |
| Participant 20 | 50 | male | Yes | Yes | Yes | 513.2 | 132.7 | 3 | 296.3 | 140 | 3 |
| Participant 21 | 50 | female | Yes | Yes | Yes | 357 | 146.5 | 3 | 611.9 | 166.6 | 3 |
| Participant 22 | 51 | male | Yes | Yes | Yes | 1092.7 | 464 | 2 | 693.2 | 126 | 4 |
| Participant 23 | 52 | male | No | Yes | Yes | 770.1 | 256.2 | 3 | 535.3 | 133.1 | 5 |
| Participant 24 | 52 | female | No | Yes | Yes | 1670.3 | 245.9 | 4 | 221.4 | 121.4 | 2 |
| Participant 25 | 52 | female | No | Yes | No | 203.2 | 164.8 | 3 | 173.5 | 88.7 | 2 |
| Participant 26 | 52 | female | Yes | Yes | Yes | 1972.6 | 344.8 | 3 | 440.6 | 204.8 | 3 |
| Participant 27 | 54 | female | Yes | Yes | Yes | 732.5 | 296.5 | 5 | 471 | 156.9 | 4 |
| Participant 28 | 55 | male | Yes | Yes | No | 632.1 | 282 | 3 | 1215.8 | 205 | 6 |
| Participant 29 | 68 | male | Yes | Yes | Yes | 937.4 | 295 | 2 | 1255.6 | 319 | 8 |
| Participant 30 | 68 | male | Yes | Yes | Yes | 619.2 | 222.5 | 4 | 1389.1 | 257.3 | 6 |
| Participant 31 | 78 | female | Yes | Yes | Yes | 450.9 | 203.6 | 5 | 632.4 | 232 | 4 |
| Participant 32 | 80 | female | Yes | Yes | Yes | 639.5 | 225.2 | 4 | 1349.2 | 249.8 | 5 |
| Participant 33 | 111 | male | Yes | Yes | Yes | 1167.5 | 284.1 | 4 | 897.7 | 185.2 | 3 |
| Participant 34 | 118 | male | Yes | Yes | Yes | 1229.3 | 332.1 | 4 | 233.1 | 145.5 | 2 |
| Participant 35 | 130 | male | No | No | No | 449.9 | 152.8 | 2 | 607.9 | 130.2 | 5 |

**Table S2.** Channel-frequency (power) and channel-timescale (complexity) clusters isolated using a stricter threshold. For the fused clusters in Table 1 (Cluster 1, 2, and 5), we isolated each oscillatory or complexity change using a stricter threshold (p = 0.0005) for clustering t-statistics. The resulting new clusters are named according to their parent cluster in Table 1 (e.g., Cluster 1a and Cluster 1b are both encompassed by Cluster 1 in Table 1). P-values are derived from empirical cluster size distributions using permutation tests. Note that we performed new permutation cluster statistics here with a stricter threshold to learn more about previously identified parent clusters rather than to test new hypotheses. Thus, we did not assess statistical significance of new clusters; however, the original parent clusters were all statistically significant (see Table 1). Effect sizes are reported as Cohen’s d (median and standard deviation across all cluster points).

|  | EEG measure | Comparison | Direction (awake-sleep) | Regressed covariates? | p-value | Cohen's d (cluster median) | Cohen's d (cluster SD) | Cluster size | percentage | Low freq | High freq | Minimum number of channels | Maximum number of channels |
| --- | --- | --- | --- | --- | --- | --- | --- | --- | --- | --- | --- | --- | --- |
| **Cluster 1a** | gMLZ | Full | Increase | No | $< 10^{-4}$ | 0.96 | 0.27 | 175 | 46.05 | 1 | 6.9 | 1 | 19 |
| **Cluster 1b** | gMLZ | Full | Increase | No | 0.0004 | 0.53 | 0.14 | 77 | 20.26 | 11.8 | 28.6 | 8 | 18 |
| **Cluster 2a** | power | Full | Decrease | No | $< 10^{-4}$ | -0.74 | 0.25 | 228 | 29.27 | 1 | 4 | 1 | 19 |
| **Cluster 2b** | power | Full | Decrease | No | $< 10^{-4}$ | -0.41 | 0.12 | 140 | 17.97 | 5.7 | 19 | 1 | 15 |
| **Cluster 5a** | gMLZ | Targeted | Increase | No | 0.0001 | 0.98 | 0.20 | 138 | 36.32 | 1 | 3.9 | 3 | 19 |
| **Cluster 5b** | gMLZ | Targeted | Increase | No | 0.0013 | 0.50 | 0.08 | 34 | 8.95 | 15.4 | 28.6 | 2 | 13 |

